# Cadmium is detrimental to *Caenorhabditis elegans* via a network involving circRNA, lncRNA and phosphorylated protein

**DOI:** 10.1101/2022.04.16.486470

**Authors:** Zhi Qu, Peisen Guo, Shanqing Zheng, Zengli Yu, Limin Liu, Panpan Wang, Fengjiao Zheng, Guimiao Lin, Peixi Wang, Nan Liu

## Abstract

Cadmium (Cd) as a heavy metal causes serious environmental pollution and multiple organ and system damage in human. However, little is known about the specific molecular mechanisms of the associated regulatory networks. In this study, we selected *Caenorhabditis elegans* (*C. elegans*) to investigate the effects of Cd exposure as it acts as an acknowledged and established genetic model organism. A total of 26 differentially-expressed circular RNA (DEcircRNAs), 143 lncRNAs (DElncRNAs), 69 microRNAs (DEmiRNAs) and 6209 mRNAs (DEmRNAs) were found and identified, which might influence reproductive function, aging processes and nervous system functions through regulating the levels of circRNAs and lncRNAs and the controlling of regulatory networks of circRNA/lncRNA-miRNA-mRNA. Based on quantitative PCR, four DEcircRNAs and three DElncRNAs were confirmed to have different expression levels between the Cd-treated and control group. Further, 5 protein-coding genes might be regulated by DElnRNAs through cis-acting and 114 by trans-acting elements. Additionally, 42 differentially regulative phosphopeptides were detected and 4 novel pairs of transcription factors (TFs)-kinase-substrate that might be influenced by Cd exposure were constructed by phosphoproteomics. Our findings suggest that Cd might influence multi-functions and the aging process of *C. elegans* and may inhibit the expression of TFs to reduce phosphorylated levels of the corresponding protein.

**Synopsis:** Cadmium exists widely in soil, water and air. This study manifested the regulatory network involving circRNA, lncRNA and phosphorylated protein in C.elegans after Cd exposure, which revealing the potential molecular mechanism underlying the toxic effect caused by Cd.

**Graphic Abstract:** 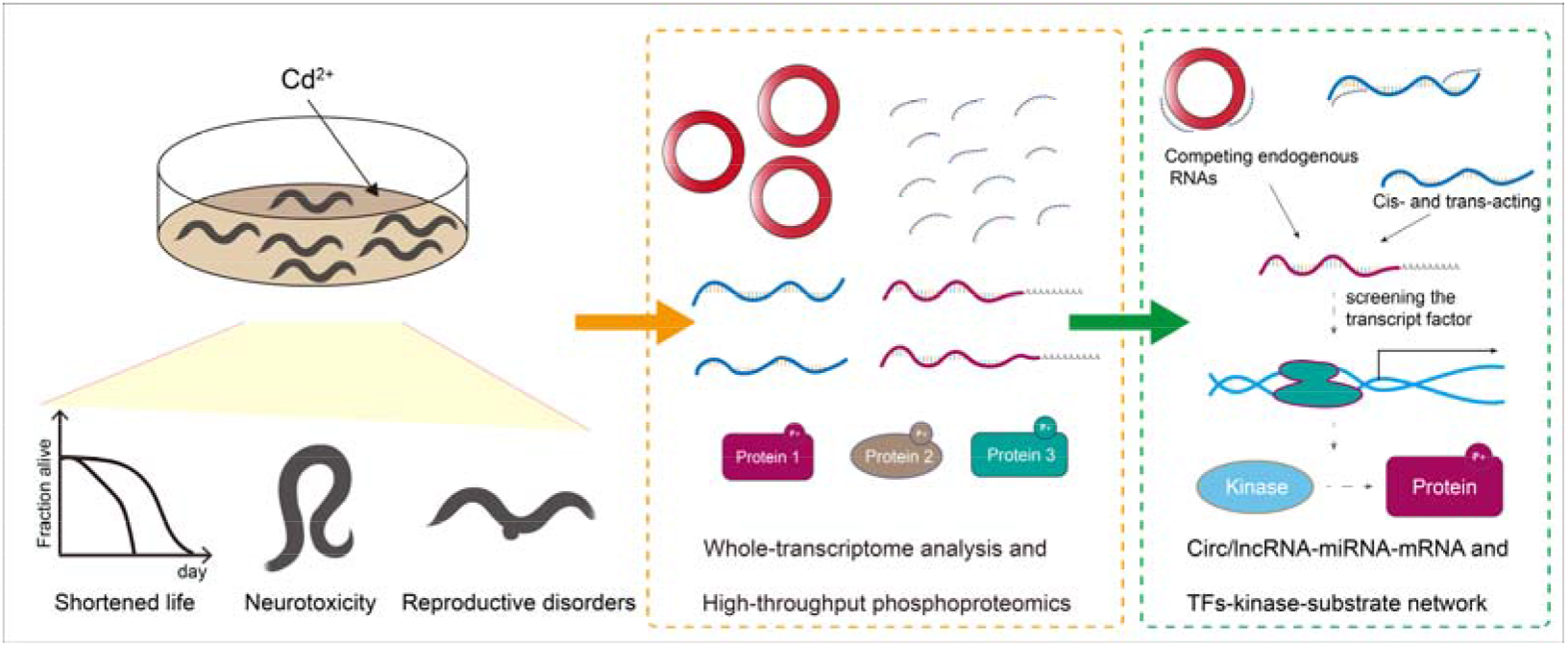

## 1. Introduction

Cadmium (Cd), as a toxic heavy metal that has adverse impact on both environmental and human health and can lead to chronic and acute diseases, such as Itai-itai and kidney disease ^1^. Due to polluted systems, especially agricultural soil, Cd can become concentrated in crops such as rice and maize ^2-4^. Cd that is absorbed through diet can accumulate in the body, especially in kidney and bones through a chronic intake process ^1, 5^. Cd exposure can cause detrimental effects, including cognitive impairment and type 2 diabetes mellitus ^6, 7^. Moreover, *in-vivo* and *in-vitro* experiments have shown that Cd is a mutagen that inhibits DNA repair ^8^ and reproductive function; experiments in rat testes showed inhibition by PI3K with the mammalian target of rapamycin (mTOR) signaling pathway, which is associated with autophagy and apoptosis ^9^. Notably, females were more vulnerable to Cd exposure than males ^10^, which would increase the risk of breast cancer (Dietary intake and urinary level of cadmium and breast cancer risk: A meta-analysis).

*Caenorhabditis elegans* (*C. elegans*) has been recognized as an appropriate model for genetic research ^11, 12^. It is considered as an ideal model organism due to work by Sydney Brenner in 1970s and possesses many advantages such as a short lifespan, simple survival condition, convenient observation, and high reproductive potential ^13^. During hermaphrodite development, if newly hatched L1 worms have no available food, the worms will enter a quiescence state called L1 arrest ^14^. During arrest, worms do not undergo postembryonic development, including cell division and growth. As such, L1-arrested worms provide an important model to study the functions of drugs and chemicals on the genetic circuits regulating cell proliferation and development. Many significant findings, including programed cell death (e.g., *ced* family and gene silencing) are based on *C. elegans* ^15, 16^. Importantly, the nervous and reproductive system of *C. elegans* is damaged by heavy metals, such as lead and Cd ^17, 18^.

Long non-coding RNA (lncRNA) has been shown to regulate cellular mechanisms, such as gene expression, histone modifications, DNA methylation, and chromatin remodeling mainly via interacting with mRNA or proteins ^19, 20^. This has been observed in many species, including *C. elegans* ^19, 21^. Based on sequence-specific binding, lncRNA can sponge the miRNA and directly interact with mRNA, commonly known as *trans-acting*, to regulate the gene expression. This process can also influence neighboring protein-coding genes by *cis-acting* ^22^.

Circular RNAs (circRNAs) were long thought to be inactive by-products due to splicing errors, and considered unlikely to play roles in biological processes (BPs). However, due to high-throughput sequencing technology, circRNAs have since been abundantly identified in eukaryotic cells. This technology has been rapidly developed with cost now declining and the methods extensively utilized in clinical and biological research ^23-25^. Furthermore, circRNAs were characterized as a function of sponge function to suppress miRNA, which a host of researchers were focused on to identify a role in many diseases, such as cancer, diabetes, and cardiovascular disease ^26^.

Many species have been sequenced using wide-genome technology and stored in a public database, including *C. elegans* ^27^. A number of scholar applied transcriptome sequencing techniques when using nematodes for their studies. Importantly, researchers have detected that circRNA in *C. elegans* ^28^ and expression level of circRNA increased with age ^29^ based on this technique, which implies that the circRNA might regulate physiological processes of *C. elegans*.

Phosphoproteomic technology has also been applied in identifying and quantifying the events of protein phosphorylation in *C. elegans* influenced by Cd. Phosphorylation is an important post-translational modification (PTM) that dynamically and reversibly regulates almost all BPs in cell. Dysregulation of phosphorylation signaling pathways is a landmark event in many pathological processes. For example, a phosphorylated protein might regulate the lifespan of *C. elegans* at different temperature and enhance the ability to resist heat and stress^30, 31^. With the development of proteomic technology based on mass spectrometry (MS), high-throughput quantitative technology and phosphorylated peptide enrichment, phosphorylated proteomics has become a mature direction of PTM. TiO_2_ possesses high specificity to phosphorylated peptides, and is widely employed in phosphorylation enrichment of serine, threonine and tyrosine.

While Cd has been confirmed to exert toxic effects on organisms including *C. elegans*; the mechanisms of how Cd exposure impacts *C. elegans* through circRNA, lncRNA and phosphorylated protein remain unclear. Here, based on transcriptome sequencing and phosphoproteomic technology, we explore the regulatory network based on circRNA and lncRNA, and the potential function influenced by exposure to Cd, which provides new insight to further investigate the mechanism of Cd toxicity at different research levels.

## 2. Material and Methods

The overall flow chart was shown in **Figure 1**.

**Figure 1.**
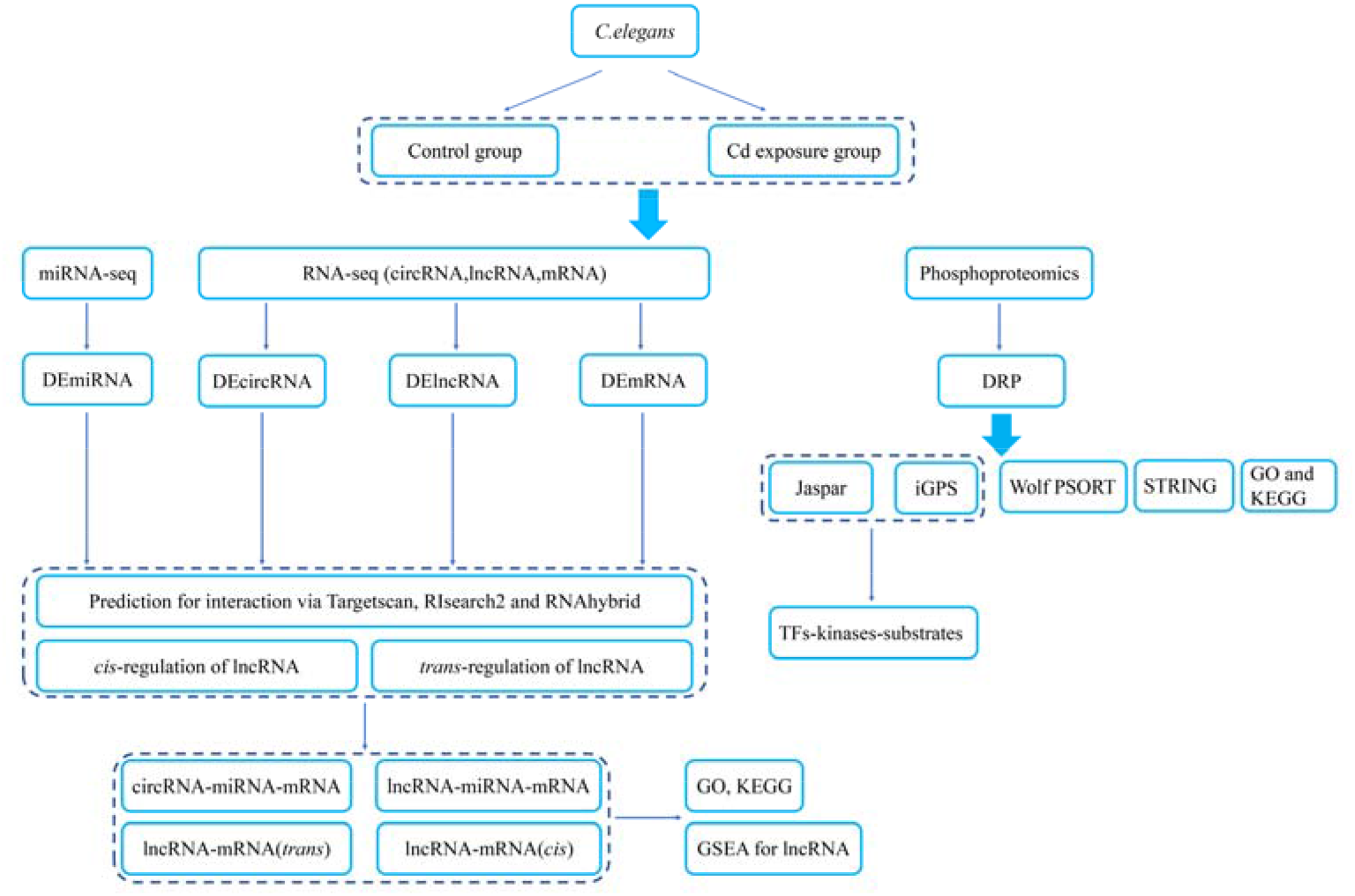
The overall flow chart of this paper.

### 2.1. Strains and media

*C. elegans* wild-type strain N2 were cultured normally on nematode growth medium (NGM) containing a food source of *E. coli* OP50 ^32^. Briefly, to collect the enough synchronized L1-larvae, the adult worms were treated with lysis solution (10% bleach and 1 M NaOH) for 5 min to release eggs. Eggs were then synchronized for 14 h to L1 stage in M9 buffer without food. L1-larvae were transferred to fresh NGM plates and incubated for 2 d at 20°C to prepare L4-larval stage worms. All strains of *C. elegans* were assessed in triplicate; all the trials were performed in triplicate.

### 2.2. Cd exposure

The method and dose of Cd for nematode toxicity were performed as previously described ^33^ with little modification. Briefly, CdCl_2_ was dissolved into 1000 μM stock solution and diluted to 60 _μ_M; L4-larvae of *C. elegans* was exposed to 60 _μ_M CdCl_2_ mixed with NGM and then added to *E. coli* with colony forming units (CFUs) of 4 × 10^6^ at 20 °C. Dead worms were removed every 24 h under a stereomicroscope. Surviving *C. elegans* were transferred to fresh NGM culture dishes by every two days; nematodes that did not respond to the stimulation with probes were defined as dead. Lifespan was employed as endpoints to evaluate the toxicity after exposure.

### 2.3. Library construction and sequencing for mRNA and circRNA

1∼2 μg total RNA of *C. elegans* was extracted from each sample and prepared for library construction. Integrity of total RNA was determined by agarose gel electrophoresis; quantification and quality control were performed by NanoDrop^®^ (ND-1000, Thermo-Fisher, USA). Total RNA was first enriched by NEB Next^®^ Poly(A) mRNA Magnetic Isolation Module (New England Biolabs, Netherlands); the library was constructed by using KAPA stranded RNA-Seq library prep kit (Illumina, USA). Bioanalyzer (G2938C, Agilent, USA) was used to control quality. The library was degraded to single-strand DNA (ssDNA) by 0.1mol/L NaOH and diluted to 8 pmol/L. Subsequently, *in situ* amplification was performed by NovaSeq 6000 S4 reagent kit (300 cycles) (15057934, illumina, USA). The product was sequenced by Illumina NovaSeq 6000.

### 2.4. Library construction and sequencing for miRNA

miRNA libraries were constructed using Multiplex Small RNA Library Prep Set for Illumina (E7508L, New England Biolabs, Netherlands). ssDNA was captured in Illumina flow cell and used for *in situ* amplification. According to the manufacturer’s instructions, the product was sequenced using Illumina NextSeq 500 (50 cycles). Other steps were similar to the mRNA sequencing protocol.

### 2.5. Raw sequencing data process for mRNA and circRNA

After quality control by FastQC (0.11.9) and discarding 3’ and 5’ adapter by Cutadapt (3.4) ^34^ and as well as short fragments (≤20bp), trimmed data was aligned to a reference genome (WBcel235) by Hisat2 (2.2.1) ^35^ and considered for further analysis according to quality. StringTie (2.1.5) ^36^ software was applied to estimate transcription abundance and determine the count of genes and transcripts. Fragments per kilobase of transcript per million mapped reads (FPKM) in gene and transcript level was calculated by the Ballgown R Bioconductor package ^37-39^; average FPKM of gene and transcript in treatment or control group < 0.1 was excluded. After exclusion, the FPKM value in gene level was used for Spearman correlation analysis for validation of sample concordance, for which the Spearman correlation should > 0.9, according to the standard of ENCODE (https://www.encodeproject.org). Then, count data of genes and transcripts were employed to explore the differentially expressed genes (DEGs) and differentially expressed transcripts using R package “DESeq2” ^40^, with the standards of *P* < 0.05 and |log2FC| > 0.585. According to the file of gene transfer format (Caenorhabditis_elegans.WBcel235.103.gtf, GTF file, http://www.ensembl.org/), the mRNA and lncRNA were identified. CircRNAs were identified and quantitated by Bowtie 2 software (2.4.2) ^41^ and find_circ (1.2) ^28^. Count data of circRNA were used to perform differentially expressed analysis for circRNA using the edgeR Bioconductor package with the standards of *P* < 0.05 and |log2FC| > 0.585 ^42^. Differentially expressed mRNAs (DEmRNAs), lncRNA (DElncRNAs) and circRNAs (DEcircRNAs) were then prepared for further analysis.

### 2.6. Raw sequencing data processing for miRNA

After quality control by Solexa CHASTITY and discarding 3’ adapter by Cutadapt (3.4) and short fragments (≤15bp), trimmed data was compared with a reference genome by using Bowtie 2 software and considered for further analysis according to quality. The miRDeep2 (0.1.3) ^43^ was used to quantitate miRNA expression. miRNA with a count per million value <1 was excluded and the count data was applied to conduct differentially expressed analysis using the edgeR Bioconductor package with the standards of *P* < 0.05 and |log2FC| > 0.585.

### 2.7. Real-time quantitative PCR for DElncRNA and DEcircRNA

N2 worms were washed three times in M9 buffer to remove bacteria. Total RNA was extracted using TRI reagent (T9424, Sigma-Aldrich, USA) and reverse transcribed using SuperScript™ III reverse transcriptase kit (18080-044, Invitrogen, USA). Primers were designed using Primer 5.0 (**Table 1**). qPCR was conducted using ViiA™ 7 Real-time PCR System (Applied Biosystems, USA) to confirm DEcircRNAs and DElncRNAs between the treatment and control groups. Validated circRNAs and lncRNAs were prepared for further analysis.

**Table 1.**
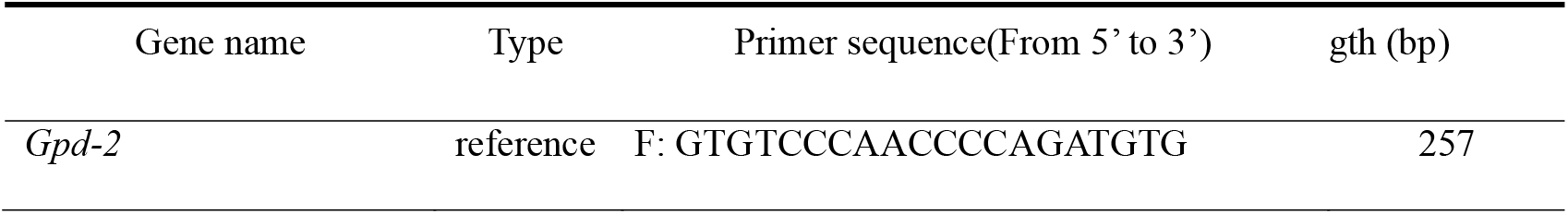
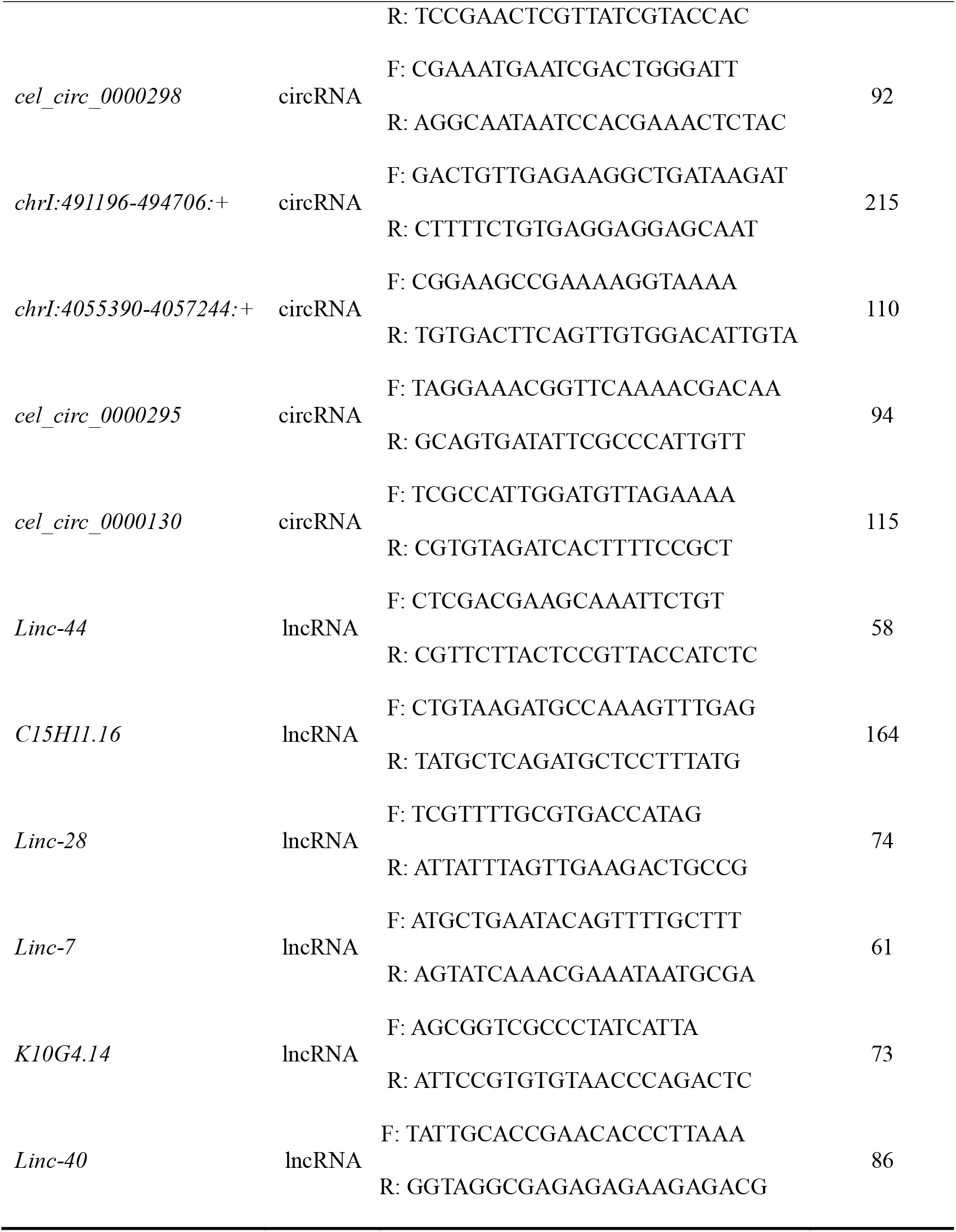
The primer of circRNAs and lncRNAs.

### 2.8. Prediction the potential target miRNAs of circRNAs and lncRNAs

The sequences of DEmiRNA were acquired from mirdeep2; and the sequences of DEcircRNA and DElncRNA were extracted according to GTF file and genome fasta file (Caenorhabditis_elegans.WBcel235.dna.toplevel.fa, http://www.ensembl.org/) by Perl language (https://www.perl.org/). Fasta sequence was produced by the Biostring Bioconductor package. The algorithms RNAhybrid ^44^, RIsearch2 ^45^ and Targetscan were employed to analyze the binding sites between miRNA and circRNA, as well as lncRNA; the results indicate that circRNA and lncRNA might have combined relationship with miRNAs. Target miRNAs were prepared for further analysis.

### 2.9. Prediction the potential target mRNA of miRNA

The seed region of miRNA typically binds with 3 prime untranslated regions (3’UTR) of the target transcript. Similar to the abovementioned methods, DEmRNA 3’UTRs were extracted using Perl. Previously noted tools including RNAhybrid, Risearch2 and Targetscan were applied to identify the miRNA targets.

### 2.10. Construction of the regulatory network of circRNA-miRNA-mRNA

The different regulatory components of these three RNA molecules were considered as targets. According to the above predictive results, the regulatory networks of circRNA-miRNA-mRNA and lncRNA-miRNA-mRNA were constructed by Cytoscape (3.8.2).

### 2.11. Gene ontology (GO) enrichment analysis and kyoto encyclopedia of genes and genomes (KEGG)

GO enrichment analysis was performed and visualized by using the Bioconductor clusterProfiler R package (4.2.2). KEGG was completed in KOBAS (2.0) (http://kobas.cbi.pku.edu.cn/). The chord chart was used to present specific genes in the enriched GO terms and drawn by GOplot.

### 2.12. Cis- and trans-regulation and gene set enrichment analysis (GSEA) on lncRNA

Gene expression might be directly regulated by lncRNA through cis- and trans-acting elements. For cis-regulation, DEmRNA 20 kb-upstream or downstream of lncRNAs were screened by Perl, and co-expression analysis with lncRNAs were conducted. Similarly, for trans-regulation, co-expression analysis was performed between DEmRNAs and lncRNAs, with the standards of *P* < 0.05 and |r| > 0.99. Next, we investigated whether BPs would be influenced by qPCR validated DEInRNAs through GSEA. The correlation between certain lncRNAs and all protein-coding genes were utilized to rank the genes. The clusterProfiler R package was used to conduct the analysis and visualized by the Bioconductor enrichplot package (1.14.2).

### 2.13. Phosphoproteome preparation and MS analysis

For each sample, cell lysate was made in RIPA buffer and quantitated by BCA assay (Pierce™ BCA Protein Assay Kit), then precipitated using acetone at -20°C. The acetone-precipitated lysate was resolubilized in HEPES with 1% sodium dodecyl sulfate. Soluble proteins were hatched for 10 min at 55°C with tris (2-carboxyethyl) phosphine (Sigma-Aldrich, St. Louis, MO, USA), then alkylated for 15 min at RT in the dark with 10 mg/mL iodoacetamide (Sigma-Aldrich, St. Louis, MO, USA). The protein mixture was then digested overnight with trypsin. Peptide mixtures were labeled by TMT10-plex Isobaric Label Reagent Set (Thermo Fisher Scientific Inc., Rockford, USA) and mixed in equal amounts. Trifluoroacetic acid (TFA) was added to precipitate sodium deoxycholate (SDC) and TMT-labeled peptides were obtained.

Next, the labeled peptides were desalted with a C18 column (St. Louis, MO, USA) and vacuum-dried. They were separated into 120 fractions using high-performance liquid chromatography (HPLC, XBridge BEH C18 XP Column, Waters, USA). Collected peptides were recombined into eight fractions according to the order and vacuum dried. Phosphopeptide enrichment was performed with TiO_2_ beads (GL Sciences Inc., Japan). For each fraction, 1/2 peptide were separated and analyzed with a Nano-HPLC (EASY-nLC1200, Thermo Fisher Scientific, Inc., USA) coupled to Q-Exactive MS (Thermo-Fisher, Finnigan). Data dependent acquisition was performed in profile and positive mode with Orbitrap analyzer at a resolution of 70, 000 (200 m/z) and m/z range of 350 -1600 for MS1; for MS2, the resolution was set at 35, 000 (200 m/z) with a fixed first mass of 110 m/z. For MS/MS, normalized collision energy was set at 32%, dynamic exclusion time was set at 30 s, automatic gain controls were set at 3E6 in MS1 and 1E5 in MS2, and the maximum ion implantation time was 50 ms (MS1) and 100 ms (MS2).

### 2.14. Phosphopeptide identification and data analysis

Raw LC-MS/MS data were processed using Maxquant (v.1.6.1.0). Tandem mass spectra were searched against the UniProt *C. elegans* database. TMT-10 plex was selected as quantification method. Trypsin/P was specified as cleavage enzyme allowing up to 4 missing cleavages. Variable modifications included oxidation (methionine), phosphorylation (serine, threonine and tyrosine) and acetylation (protein N-terminal). The fixed modification included Carbamidomethylation (cysteine). False discovery rate (FDR) thresholds for protein, peptide and modification site were specified at 1%. Minimum peptide length was set at 7. Number of modifications per peptide was not greater than 5. Maximum molecular mass of per peptide was 4, 600 Da. All the other parameters in MaxQuant were set as default values, and only three types of phosphorylation including serine, threonine and tyrosine were quantified.

For data analysis, t-test was used to identify the differentially regulative phosphopeptides (DRPs) with the criteria of |log_2_FC| ≥ 0.585 and *p*-value < 0.05); fold change (FC) was calculated through the ratio of treatment to control group based on the average quantity of phosphorylated peptide. Functional classifications of phosphoproteins were conducted using the Bioconductor clusterProfiler R package. Subcellular locations were predicted using online tool Wolf PSORT (https://wolfpsort.hgc.jp/) ^46^. Protein-protein interactions (PPI) network was detected using STRING database ^47^. The corresponding kinase of per DEPs was identified using iGPS software ^48^. Then, the Perl was used to extract the upstream 200 bp of these genes as promoter sequence, which was submitted to Jaspar database (http://jaspar.genereg.net/) to predict the corresponding transcription factors. Based on the above RNA-seq data, differentially-expressed transcriptional factors were identified.

### Statistical analysis

For qRT-PCR, student’s t-test was used to identify differences between treatment and control group. For RNAhybrid, the seed region of miRNA was forced to pair with the target. To enhance accuracy, cut-off values of minimum free energy (MFE) was set as -18 for predicting the combination relationships in the pairs of lncRNA-miRNA and circRNA-miRNA, and -20 for miRNA-mRNA. For RIsearch, the standard of MFE was < -10. For all three tools, the sequence between 2nd and 8th of 5’end of miRNA was regarded as the seed region, which should pair with the target sequence.

## 3. Results

### 3.1. Quality of raw sequencing data and differentially expressed analysis

All raw sequencing data were eligible for further analysis, results of quality control for miRNA and other RNA types are shown in Supporting Information (SI) Figures S1, S2. The results of mapping to genome through Hisat2 showed a higher concordant rate (**Table 2)**. The distribution of gene expression level in each sample is shown in SI Figure S3. Spearman analysis showed the replicated concordance and good performance (**Figure 2A**). According to the standards, 26 DEcircRNAs, 69 DEmicroRNAs, 6209 DEmRNAs and 144 DElncRNAs were identified. Among these, 20 circRNAs were up-regulated and 5 were down-regulated (**Figure 2B**); 108 lncRNAs were up-regulated and 36 were down-regulated (**Figure 2C**); 3259 mRNAs were up-regulated and 2950 were down-regulated; 30 miRNAs were up-regulated and 39 were down-regulated. The difference of expression level of DEcircRNAs and DElncRNAs is shown in **Figure 2D** and **E**, which drawn by the software of TBtools (1.6) ^49^

**Table 2.** The sequencing raw and trimmed data mapping to the reference genome for each group.

| samples | Raw pairs | Trimmed | Mapped (%) |
| --- | --- | --- | --- |
| 2-1 | 65291348 | 65289765 | 79.12 |
| 2-2 | 41132724 | 41131912 | 85.72 |
| 2-3 | 44442403 | 44441700 | 92.94 |
| 2-4 | 50483810 | 50483125 | 94.27 |
| 2-5 | 45470358 | 45469436 | 90.58 |
| 2-6 | 45165010 | 45164076 | 87.21 |

**Figure 2.**
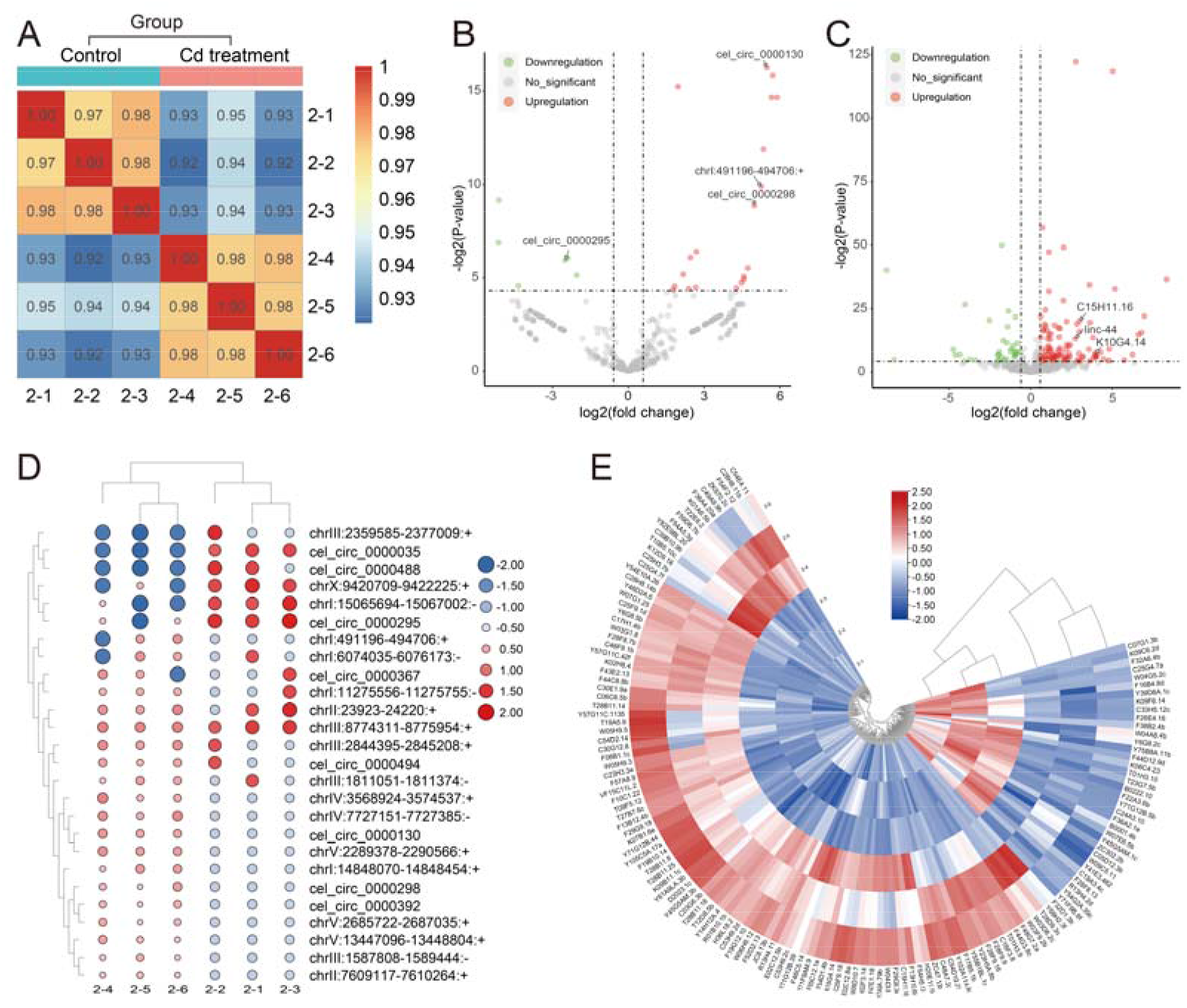
Sample concordance and differentially expressed analysis. **A**. Sample concordance between each group; **B**. Volcanic plot for circRNA; four validated circRNAs are annotated; **C**. Volcanic plot for lncRNA; three validated lncRNAs are annotated; **D**. Heatmap for DEcircRNAs; **E**. Heat map for DElncRNA.

### 3.2. Validation of circRNA and lncRNA expression and prediction of target miRNA

Five circRNAs and six lncRNAs were further analyzed by qPCR to confirm differential expression. Four circRNAs *cel_circ_0000298, chrI:491196-494706:+, cel_circ_0000295* and *cel_circ_0000130*, and three lncRNAs *Linc-44, C15H11.16* and *K10G4.14* were analyzed and it was confirmed that their expression levels were significantly different between treated and control groups (**Figure 3A**). Their target miRNAs were predicted by RNAhybrid, RIsearch2 and Targetscan; 45 pairs of circRNA-miRNA (**Figure 3B**) and 11 pairs of lncRNA-miRNA (**Figure 3C**) were found, the former included three unique circRNAs and 38 unique miRNAs (**SI Table S1**) while the latter contained three unique lncRNAs and 11 unique miRNAs (**SI Table S2**). Interestingly, all these three circRNAs and three lncRNAs were up-regulated. Therefore, 15 up-regulated miRNAs in predictive results were excluded for circRNA and five up-regulated miRNAs for lncRNA.

**Figure 3.**
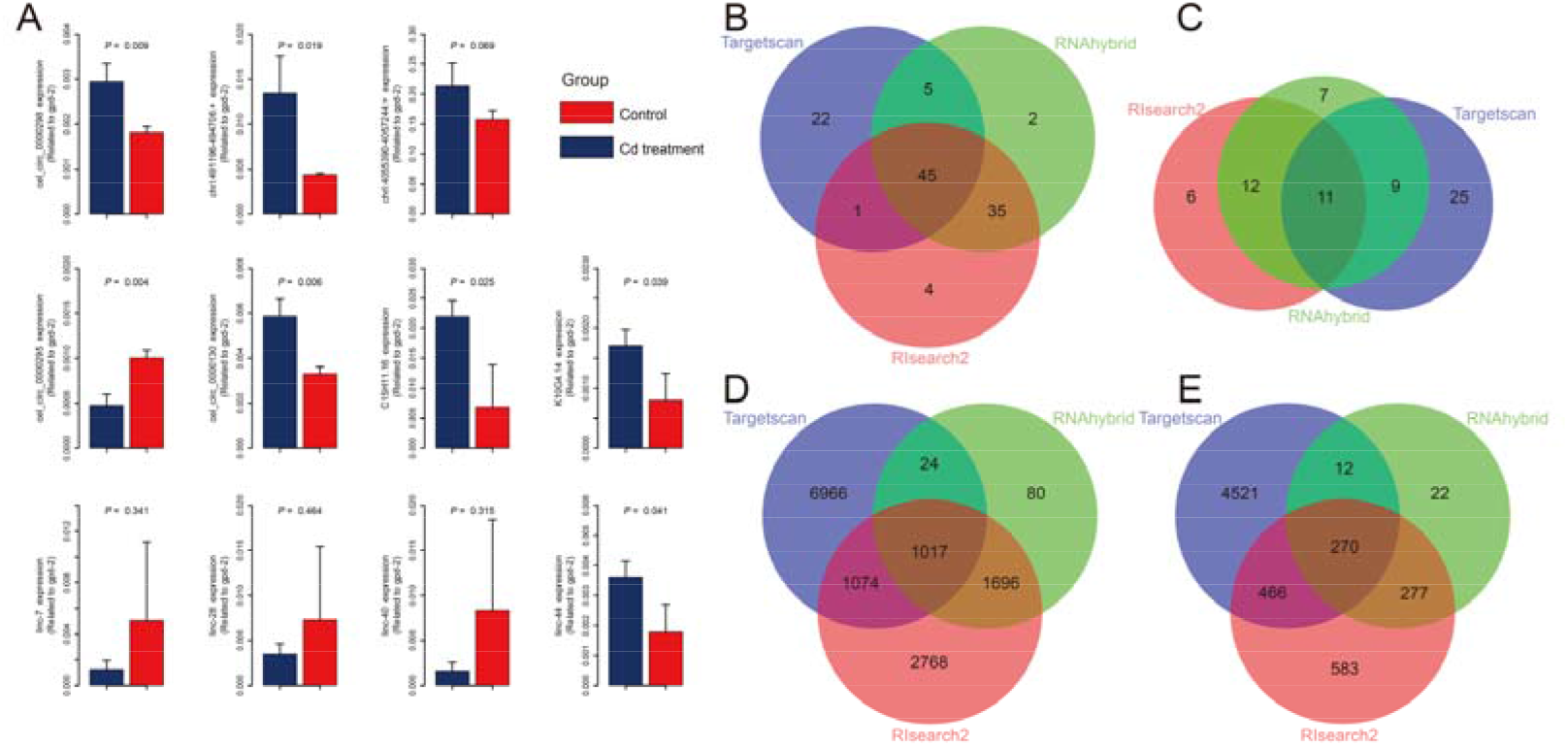
Real-time PCR and prediction of miRNA-RNA interaction. **A**. Real-time PCR assay to confirm the different expression of five DEcircRNAs andsix DElncRNAs between treatment and control groups; **B**. Prediction of the circRNA-miRNA pairs and intersection of the results of three predictive tools; **C**. Prediction of the lncRNA-miRNA pairs and intersection of the results of three predictive tools; **D and E**. Prediction of miRNA-mRNA pairs downstream of circRNA and lncRNA, respectively, and intersection of the results of three predictive tools.

### 3.3. Prediction of the target mRNA of miRNA

After identification of above-mentioned target miRNAs of circRNA and lncRNA, three tools were used to conduct the prediction. For circRNA, a total of 1017 miRNA-mRNA pairs were identified (**Figure 3D**), containing of 316 unique genes after excluding mRNAs whose direction was identical with miRNA (**SI Table S3**); for lncRNA, 270 miRNA-mRNA pairs were identified (**Figure 3E**), containing 114 unique genes after applying similar exclusion criteria (**SI Table S4**).

### 3.4. Regulatory network of circRNA-miRNA-mRNA and lncRNA-miRNA-mRNA

Based on the prediction of target miRNA and mRNA, networks of circRNA-miRNA-mRNA and lncRNA-miRNA-mRNA (**Figure 4**) were constructed by Cytoscape. GO and KEGG analysis were performed on downstream genes of circRNAs (**Figure 5A and B**). The specific genes under the enriched terms in GO analysis were displayed in a chord chart (**Figure 5C and D**). Similarly, analyses of downstream genes of lncRNA were conducted (**Figure 5E and F)**; specific genes are shown in **SI Table S5**.

**Figure 4:**
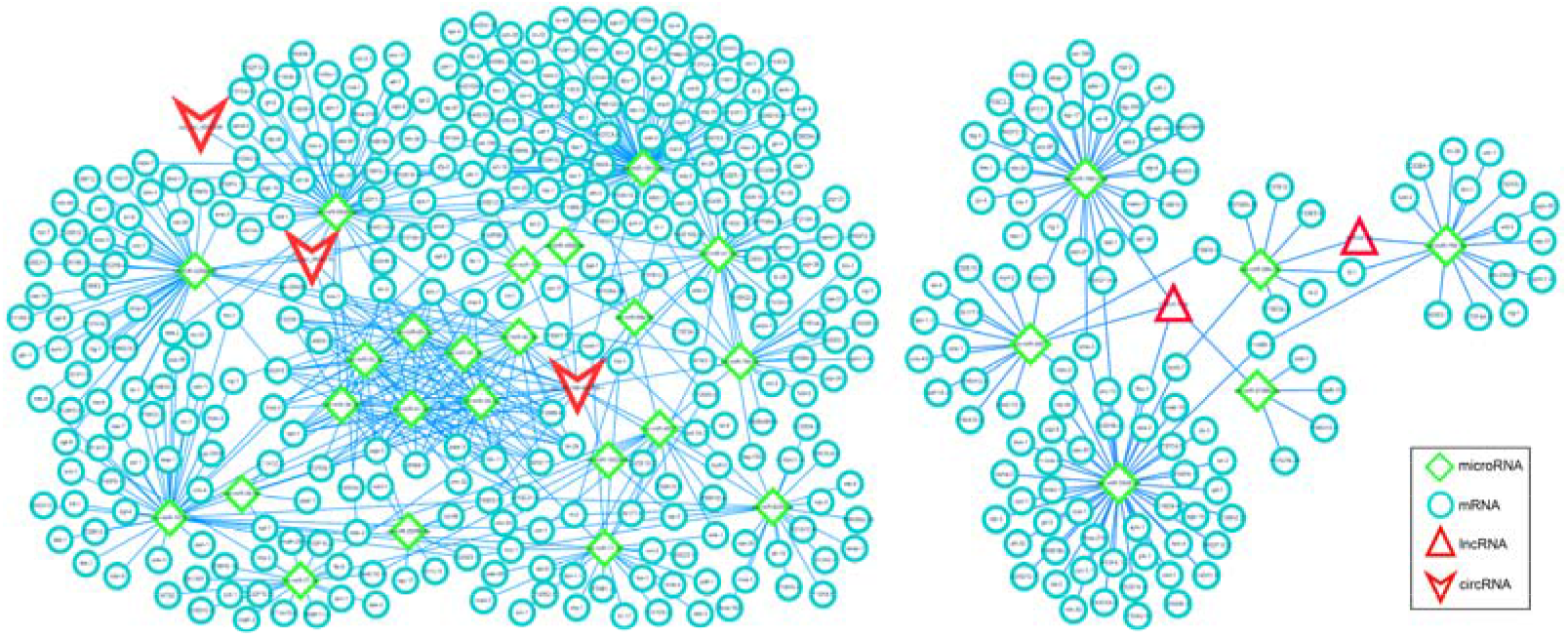
Molecular regulatory network of circRNA/lncRNA, miRNA and mRNA. Competing endogenous RNAs network was constructed by circRNA/lncRNA, microRNA and mRNA. Red V-shapes represent circRNA; green diamonds represent miRNA; red triangles represent lncRNA; annular shapes represent mRNA.

**Figure 5:**
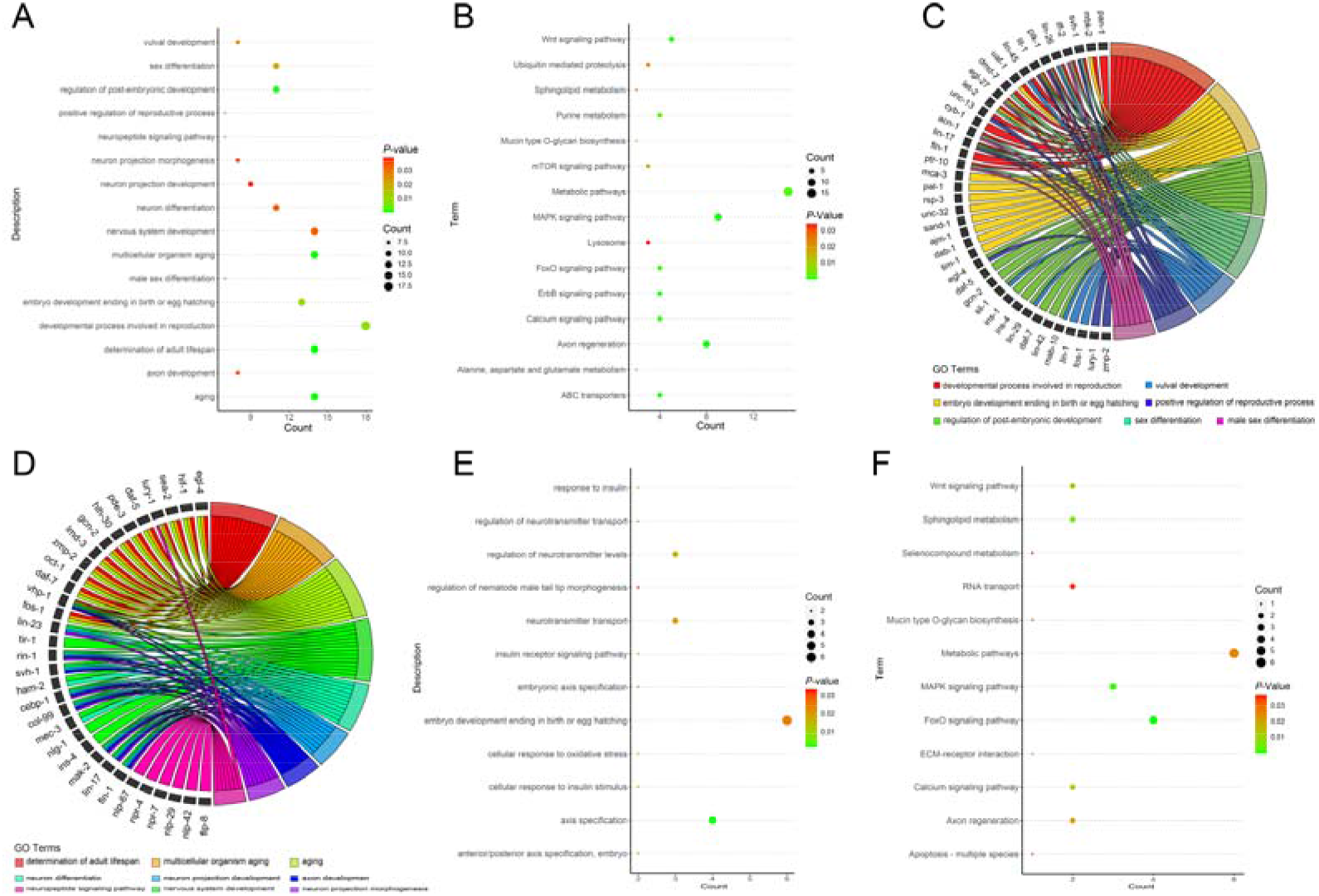
GO and KEGG analysis. **A**. GO analysis on mRNA of the circRNA downstream; **B**. KEGG analysis on mRNA of the circRNA downstream; **C**. Specific genes under the terms associated with reproduction in the GO analysis downstream of circRNA; **D**. Specific genes under the terms associated with aging and nerve in the GO analysis on the downstream of circRNA; **E**. GO analysis on mRNA of the lncRNA downstream. **F**. KEGG analysis on mRNA of the lncRNA downstream.

### 3.5. Cis- and trans-regulation of lncRNA

For cis-acting elements, five proteins encoding *Dmd-5, Acdh-8, Sri-70, K10G4.5*, and *Gnrr*, in the range of 20 kb upstream and downstream of the above-mentioned three lncRNAs were found (**Figure 6A**). According to Pearson correlation analysis, 133 DEmRNAs might be regulated by trans-acting elements (**Figure 6B**). The location of DEmRNAs, DEcircRNA and DElncRNA in chromosome and trans-acting relationship are shown in **Figure 6C**. The method of GSEA on three lncRNAs including *C15H11.16, K10G4.14, Linc-44* are exhibited in **Figure 6D-F**, respectively.

**Figure 6:**
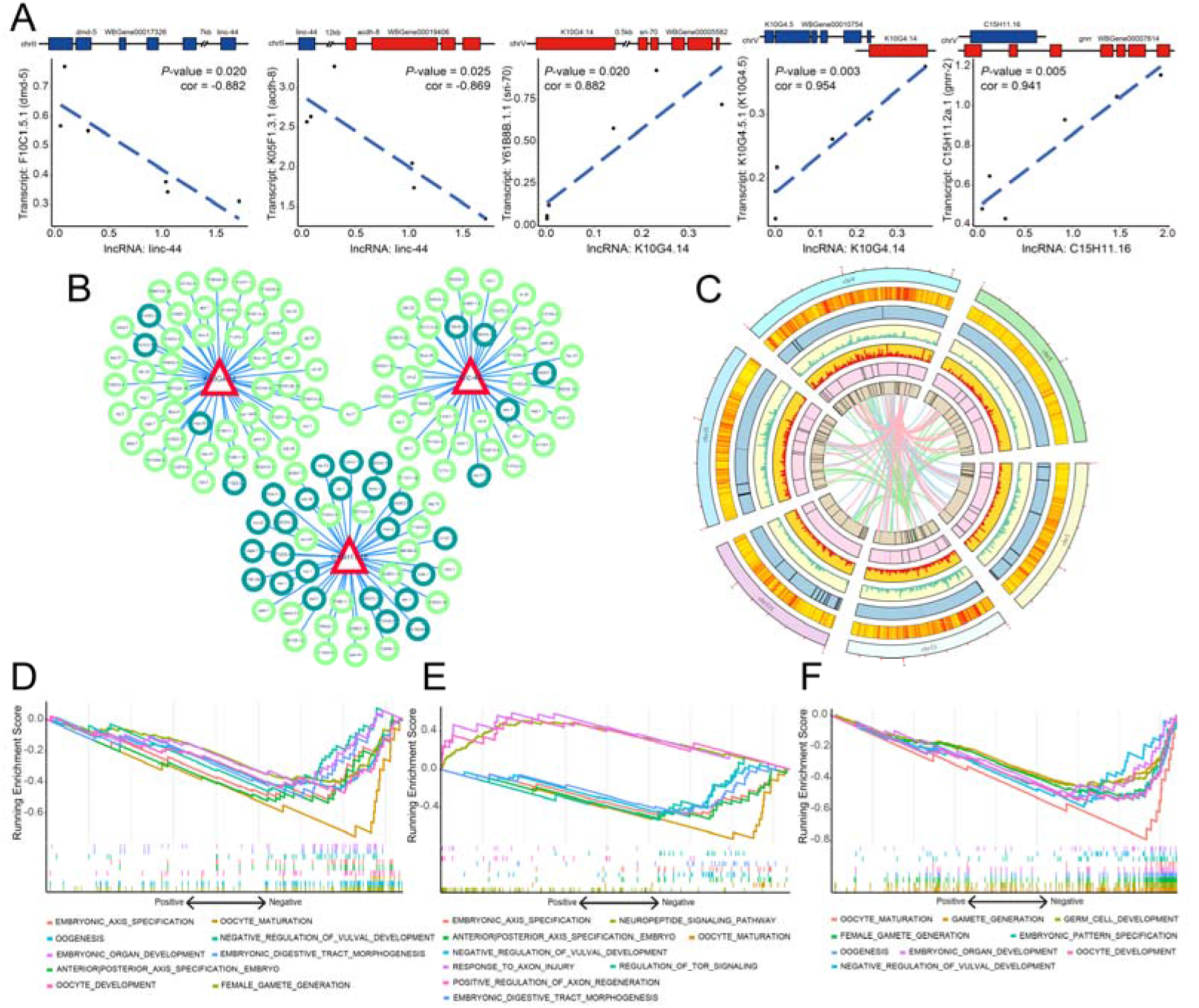
*Cis-* and *trans-acting* of lncRNA. **A**. *Cis-acting* of lncRNA on neighboring protein-coding genes; **B**. *Trans-acting* of lncRNA; **C**. The outermost layer represents the six chromosomes; the second layer represents distribution of all *C. elegans* genes; the third represents all DEcircRNAs, the fourth and fifth represent the downregulated and upregulated mRNAs, respectively; the next two layers represent downregulated and upregulated lncRNAs, respectively; the center lines represent trans-action of lncRNA on corresponding mRNAs, shown by green, pink and blue lines respectively, for *Linc-44, K10G4.14* and *C15H11.16*; **D-F**. The results of GSEA, which revealed change in function of reproduction and nerve influenced by *Linc-44, K10G4.14* and *C15H11.16*, respectively.

### 3.6. Phosphoproteomic analysis of C. elegans by Cd exposure

After normalization, a total of 200 phosphorylated peptides were eligible for further analysis. Among them, 139, 18, and 5 phosphopeptides with one, two, and at least three phosphosites, respectively (**Figure 7A**). The percentage of phosphorylated phosphosites on serine, threonine, and tyrosine were 87% (174/200), 11% (22/200), and 2% (4/200), respectively (**Figure 7B**). A total of 42 DRPs are displayed in **Figure 7C** including 15 up-regulations and 27 down-regulations. The subcellular localization of DRPs is shown in **Figure 7D**, many are nuclear. The DRPs were also be visualized using a heat map (**Figure 7E**). GO analysis of all differentially regulated phosphoproteins are shown in SI Figure S4. The relationship between each enriched term in BPs is displayed by DAG (direction acyclic graph) (**SI Figure S5**); specific terms enriched in BP and the number of genes under the corresponding terms is presented by a pie chart (**SI Figure S6**). Similarly, KEGG results of these DRPs were related to terpenoid backbone biosynthesis and autophagy (**SI Figure S7**). PPI network is shown in SI Figure S8.

**Figure 7.**
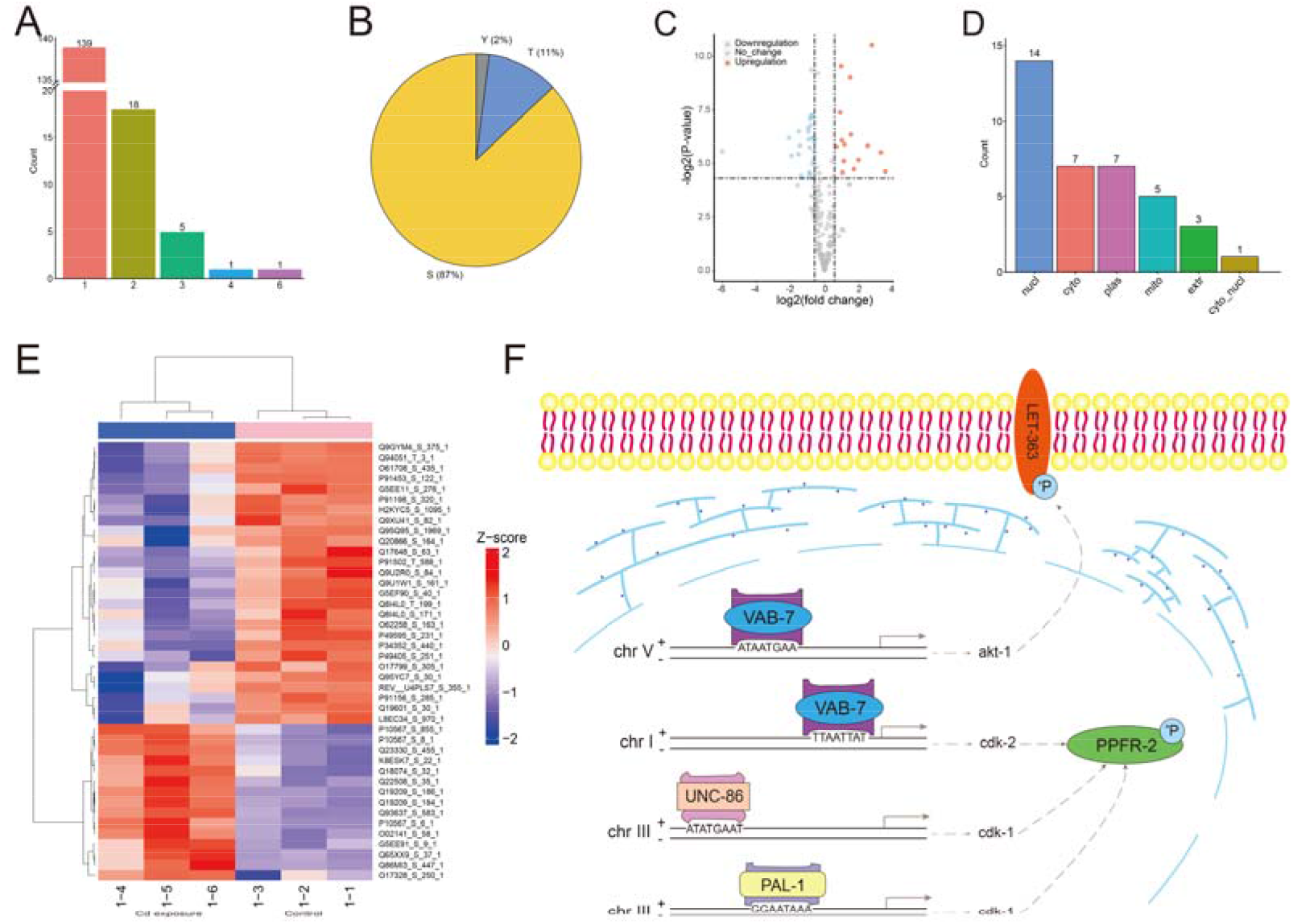
Results of phosphoproteomic data analysis. **A**. Number of phosphorylated sites for each protein; **B**. Proportion of each type of phosphorylated site (S, Y, and T); **C**. The differentially regulative peptides; **D**. Counts for location of the differentially regulative proteins; **E**. The phosphorylated peptides and relative expression in cluster heatmap; **F**. Regulative relationships of TFs (*VAB-7, UNC-86 and PAL-1*)-kinases (*akt-1, cdk-1 and cdk-2*)-substrates (phosphorylated proteins: PPFR-2 and LET-363).

In prediction of the kinase-substrate interaction pairs in *C. elegans* after Cd exposure, substrate sites were found in seven kinases, among which the regulative direction (down regulation) of three kinases (**SI Table S6**), *Cdk-1, Cdk-2* and *Akt-1*, were identical to the phosphorylated proteins. Predictive transcription factors (TFs) of these three kinases were identified and specific binding sites are shown in **SI Table S7**; three TFs, *Vab-7, Unc-86* and *Pal-1*, have the same direction (down regulation) with their corresponding gene. The predictive TFs-kinase-substrate axis is shown in **Figure 7F**.

## 4. Discussion

The present study revealed the correlative regulatory network of circRNAs and lncRNAs of *C. elegans* after Cd exposure and the related functions that are interfered with or inhibited via circRNAs and lncRNAs. Transcriptome sequencing was performed to detect differentially expressed RNA. Notably, in processing the raw sequencing data, we did not predict the novel transcripts through setting relative parameters, which could lead us to pay ample attention to analyze the known transcripts. Spearman analysis demonstrated that the relevance of intra-group samples was higher than that of inter-groups, which indicates all the replicated samples could be included in the next analysis. Then, since the sample size was smaller than the recommended number, a ballgown analysis was just used to extract the FPKM value of transcripts and genes ^39^. Unlike previous studies, we utilized qPCR to validate the expression of lncRNA and circRNA after differential expression analysis; confirmed sequences were included in further analysis, which enhanced the reliability of our analysis.

Both circRNA and lncRNA could act as sponges to inhibit miRNA and influence the expression of mRNA. Potential miRNA targets could be predicted by taking advantage of tools such as TargetScan, miRanda, RNA22 and RNAhybrid ^50^. The various algorithms used by these programs cause differences in predictive results. Therefore, to enhance accuracy, three tools were applied and relatively strict parameters were set, and we obtained their intersection. It is worth noting that when a possible binding site was predicted by RIsearch2, the positive strand in the results was eligible for predictive analytics and the minus strand was excluded. The minus-strand in the results was complementary to the sequence we input, which was different from our intention. We only need the predictive information for the positive-strand (i.e., input RNA) sequence.

GO and KEGG analyses were applied and revealed that Cd exposure might influence reproductive function, aging processes and nervous system function under the regulation of circRNA. Similarly, reproductive and nervous functions might be inhibited by lncRNA through Cd exposure. For GO analysis, we focused on biological processes. While parameter settings were strict, the obtained eligible terms were still numerous. We selected terms associated with reproduction, aging and nerve instead of the top ones, which might be important in development and were main target of Cd ^51, 52^.

Phosphorylated proteomic technology was also employed in this study, almost all phosphorylated proteins could be provided by Cd exposure in *C. elegans*. We integrated RNA-seq and phosphorylation data to explore or identify related signaling pathways. However, kinases that promote phosphorylation might not be regulated by the discovered circRNA and lncRNA under the presupposed standards. Commonly, mRNA is translated as TFs to increase the expression of a kinase, which then promotes the phosphorylation of its substrate. According to our phosphorylated proteomics and RNA-seq data, we have identified both DRPs and DEGs. Then, kinase-substrate was predicted and the differentially-expressed kinases were screened. While interactions between kinase and substrate could be regulated by other factors, kinase expression level could only be considered to influence the phosphorylation level and the expressed direction of kinase should be the same as the phosphorylated protein; the same is true for TFs and kinase expression.

Cd is known to be an environmental endocrine disruptor, which can be harmful to the reproductive system, in both males and females ^53^. Previous work has shown that Cd can produce adverse reproductive effects in mice ^54^, rats ^55^ and *C. elegans* ^56^. However, the specific molecular mechanisms remain unclear. Based on transcriptome sequencing, we identified that Cd might impact reproductive and nervous function, as well as aging processes through regulating the expression of circRNAs and lncRNAs in *C. elegans*. One limitation of this study was that the discovered circRNAs were not confirmed by Sanger sequencing ^57^ and expression level was dramatically lower than the corresponding linear transcripts and miRNA. The sample size was also limited. It would also be beneficial to treat *C. elegans* with different concentrations of Cd, in order to obtain exact dose-response relationship between gene expression and Cd. We offer the novel idea that Cd might influence on reproduction, aging and nervous system function through circRNA and lncRNA. Future works are warranted to understand the molecular significance and explore the mechanisms of circRNA, lncRNA and phosphorylated protein on gene expression. In the present study, we provide the molecular regulatory networks of circRNA/lncRNA-miRNA-mRNA using the model organism *C. elegans* after Cd exposure. This indicates a novel model and gives insight into the molecular toxicological effects of Cd. Our study revealed that the external Cd exposure might mainly influence the reproductive and nervous system function and aging through regulating the networks. Additionally, transcription factor expression might be inhibited to reduce the phosphorylated level of the corresponding protein. Future work is required to understand the molecular significance of circRNA, lncRNA and phosphorylated protein and to explore the specific mechanisms at play.

## Availability of data and materials

RNA-seq and miRNA-seq data have been submitted to the Gene Expression Omnibus (GEO accession numbers GSE189666). Source data are provided with this paper. The mass spectrometry proteomics data have been deposited to the ProteomeXchange Consortium (http://proteomecentral.proteomexchange.org) via the iProX partner repository with the dataset identifier PXD030083.

## CRediT authorship contribution statement

Zhi Qu: Methodology, Funding acquisition, Writing -review & editing. Peisen Guo: Methodology, Writing - original draft, Writing -review & editing, Visualization. Shanqing Zheng: Writing -review & editing, Funding acquisition, Project administration. Zengli Yu: Writing -review & editing, Project administration. Limin Liu: Data acquisition. Panpan Wang: Data acquisition. Fengjiao Zheng: Writing - editing, Funding acquisition. Peixi Wang: Writing - editing. Guimiao Lin: Writing - editing. Nan Liu: Methodology, Funding acquisition, Writing -review & editing, Project administration.

## Acknowledgments

We would like to acknowledge Prof. Pingchang Yang’s (Health Science Center, Shenzhen University) kindly help in academic supporting and Aksomics for assistance with RNA-sequencing and phosphoproteomics.

