## Supplemental information caption for "Cadmium is detrimental to *Caenorhabditis elegans* via a network involving circRNA, lncRNA and phosphorylated protein"

Supplementary Fig. 1 and Fig. 2 indicated that the quality of trimmed data were eligible, which the average score of base at each position were greater than 30.

Supplementary Fig. 3 shown the distribution of sequencing data was consistant with each other.

Supplementary Fig. 4 presented the GO analysis on the differentially regulative protein.

Supplementary Fig. 5 was directed acyclic graph that presented biological process affected by cadmium.

Supplementary Fig. 6 shew that the top terms and the number of enriched genes based on its P-value.

Supplementary Fig. 7 revealed the significant signal pathway affected by cadmium.

Supplementary Fig. 8 was the PPI network based on the differentially regulative protein.

Supplementary Table 1, 2 presented the predictive circRNA-miRNA pairs and lncRNA-miRNA pairs.

Supplementary Table 3 was the predictive miRNA-mRNA pairs in the downstream of circRNA.

Supplementary Table 4 was the predictive miRNA-mRNA in the downsteam of lncRNA.

Supplementary Table 5 was shown the enriched terms of biological process and its specific genes.

For the phosphorylated proteomic, the software of iGPS was used to predict the kinase-substrate relationship and the threshold was set as high. The results was shown in Supplementary Table 6.

The specific binding site between TFs and kinases predicted by Jaspar database were shown in Supplementary Table 7.
