## Supplemental figures and tables for "Cadmium is detrimental to *Caenorhabditis elegans* via a network involving circRNA, lncRNA and phosphorylated protein"

Supplementary Fig. 1 The mean quality scores of trimmed data of RNA-sequencing.

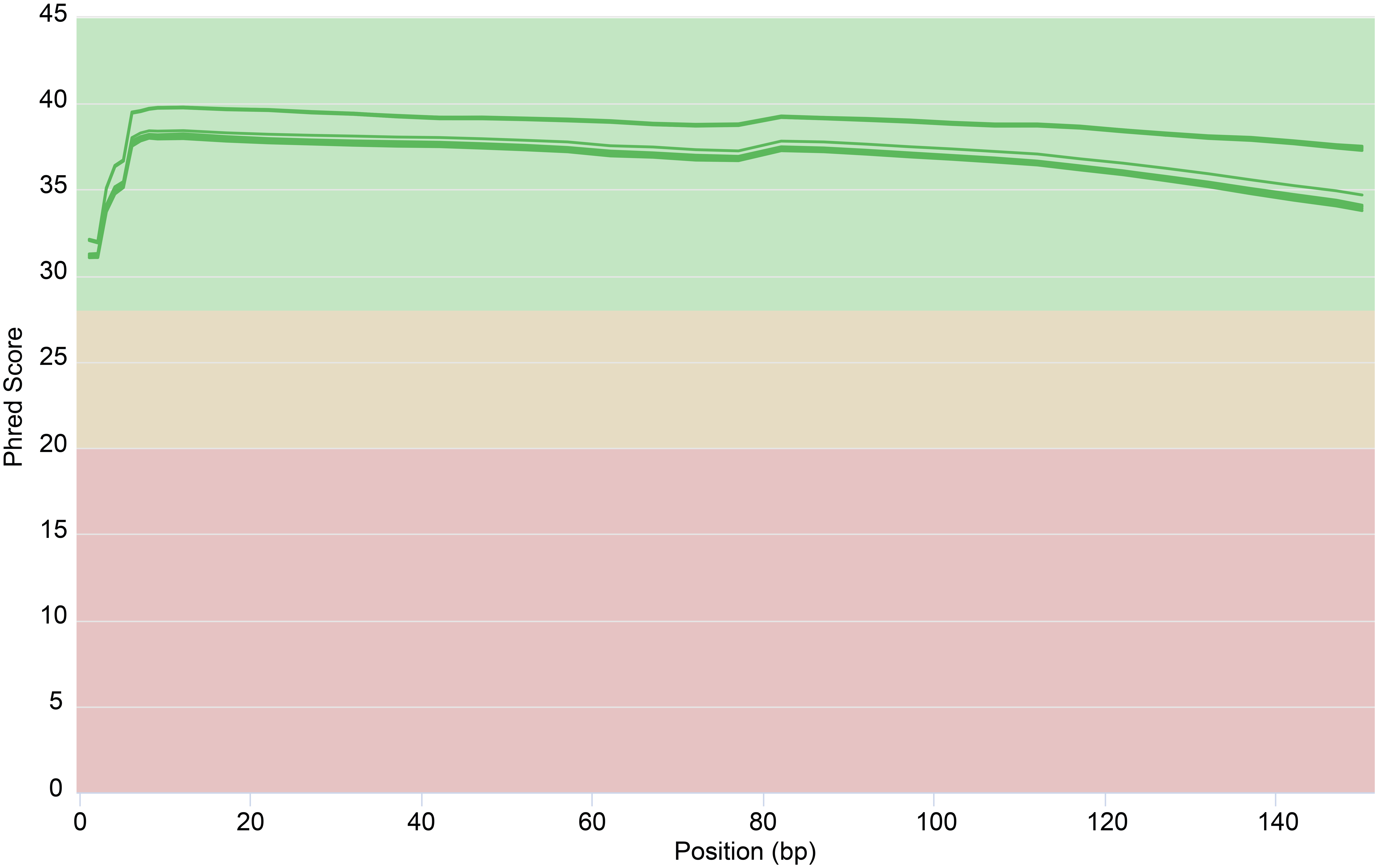

Supplementary Fig. 2 The mean quality scores of trimmed data of miRNA-sequencing.

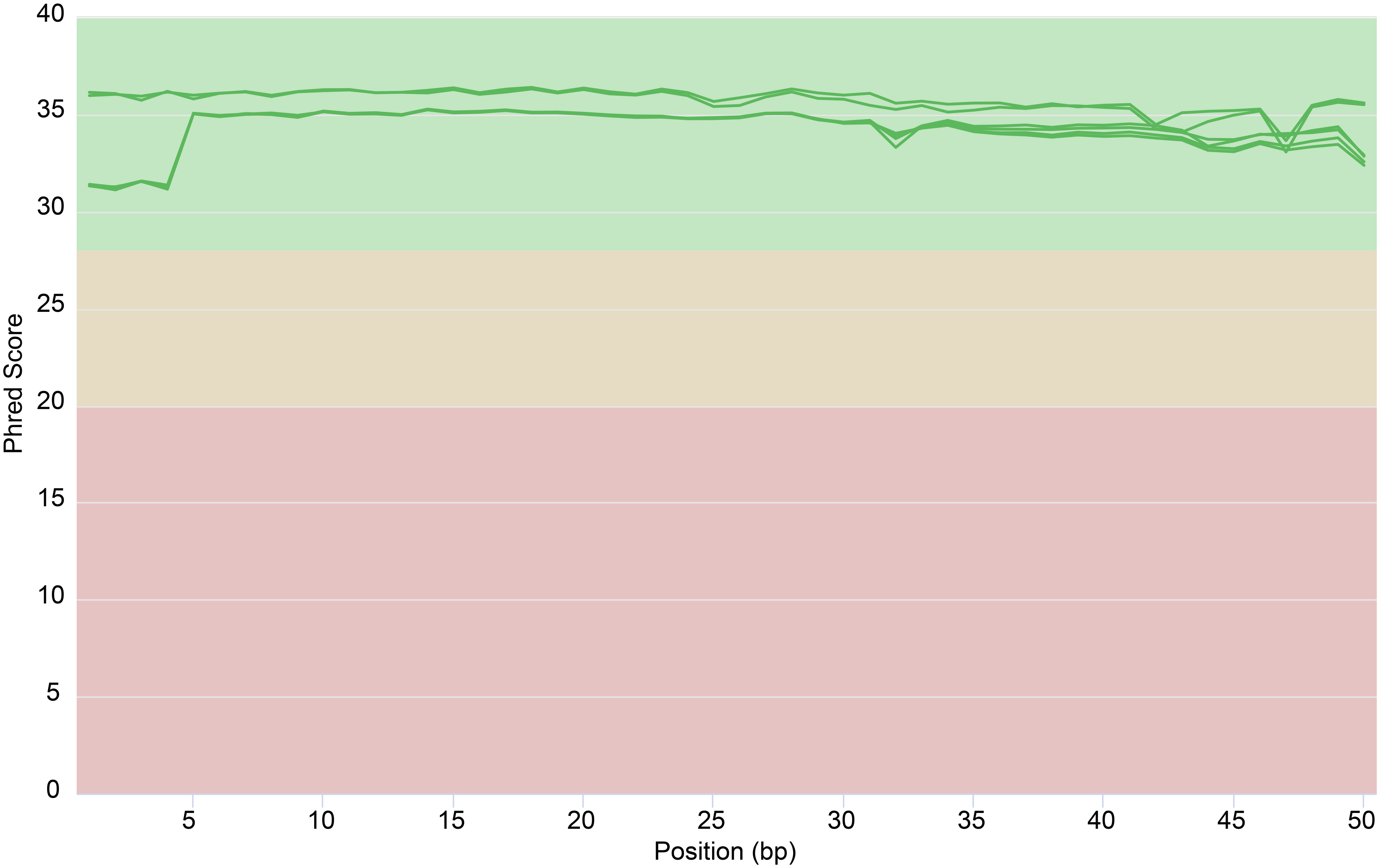

Supplementary Fig. 3 The distribution of gene expression level in six groups.

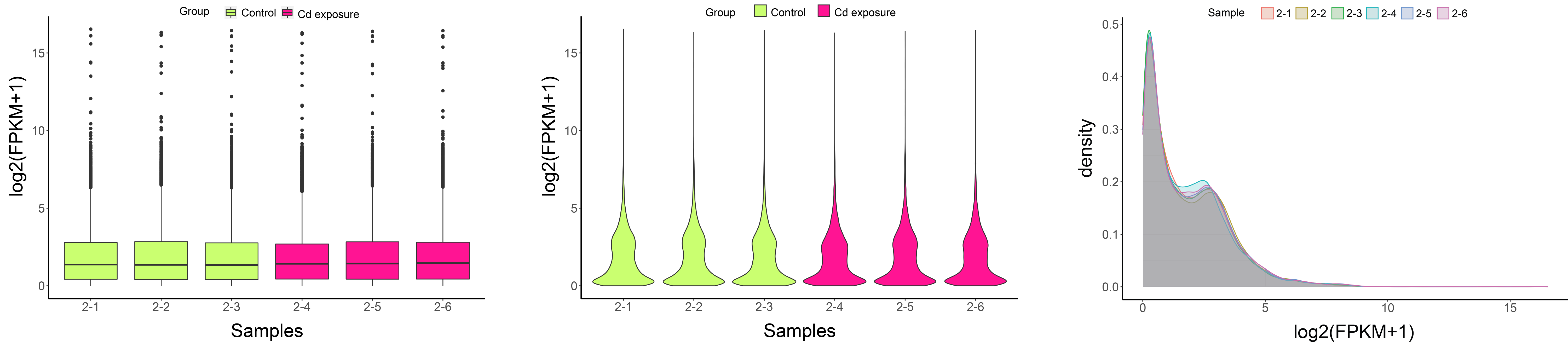

Supplementary Fig. 4 The GO analysis on differentially regulative protein

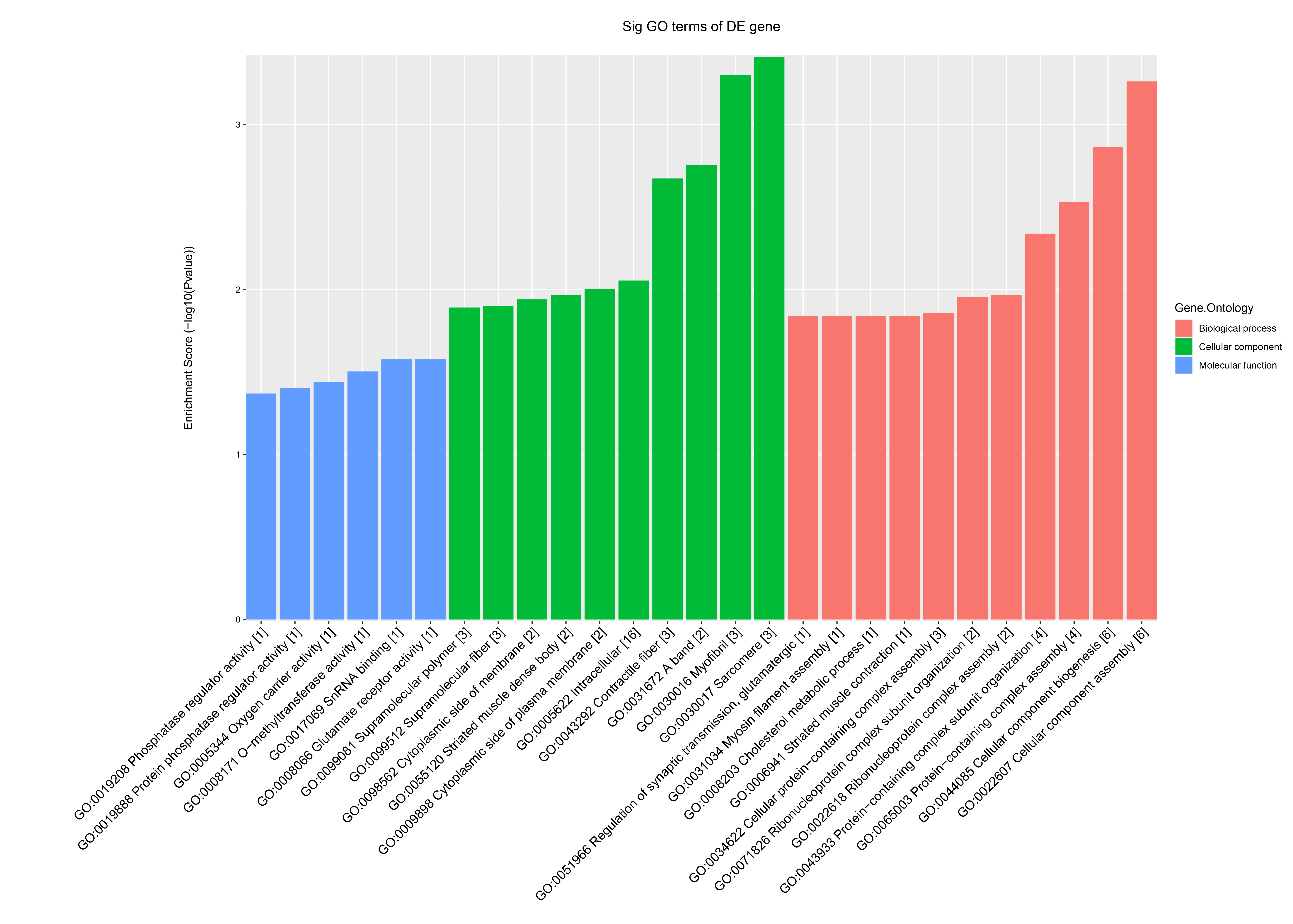

Supplementary Fig. 5 The directed acyclic graph of biological process in GO analysis.

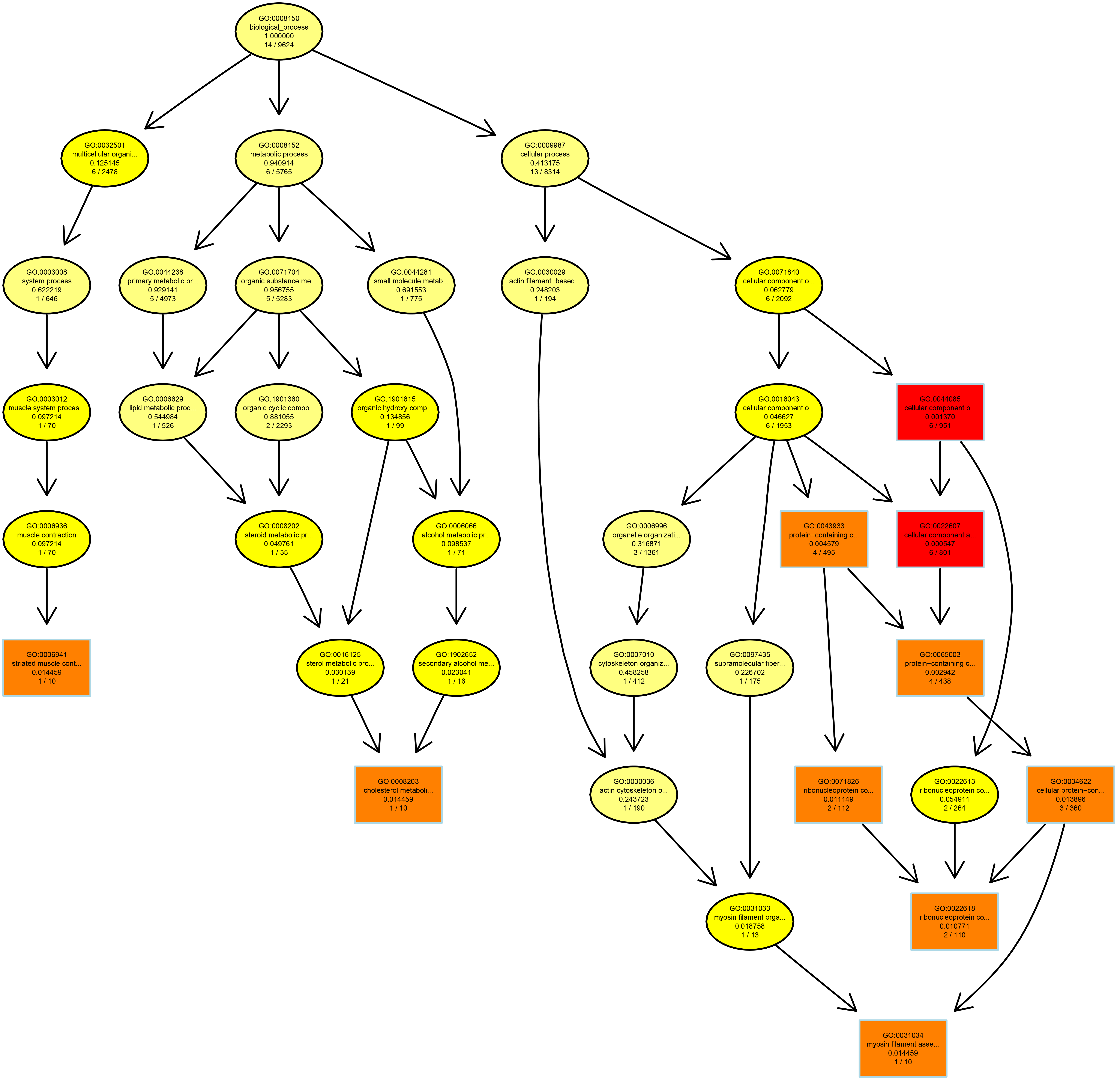

Supplementary Fig. 6 The top terms and the number of enriched genes in GO biological process.

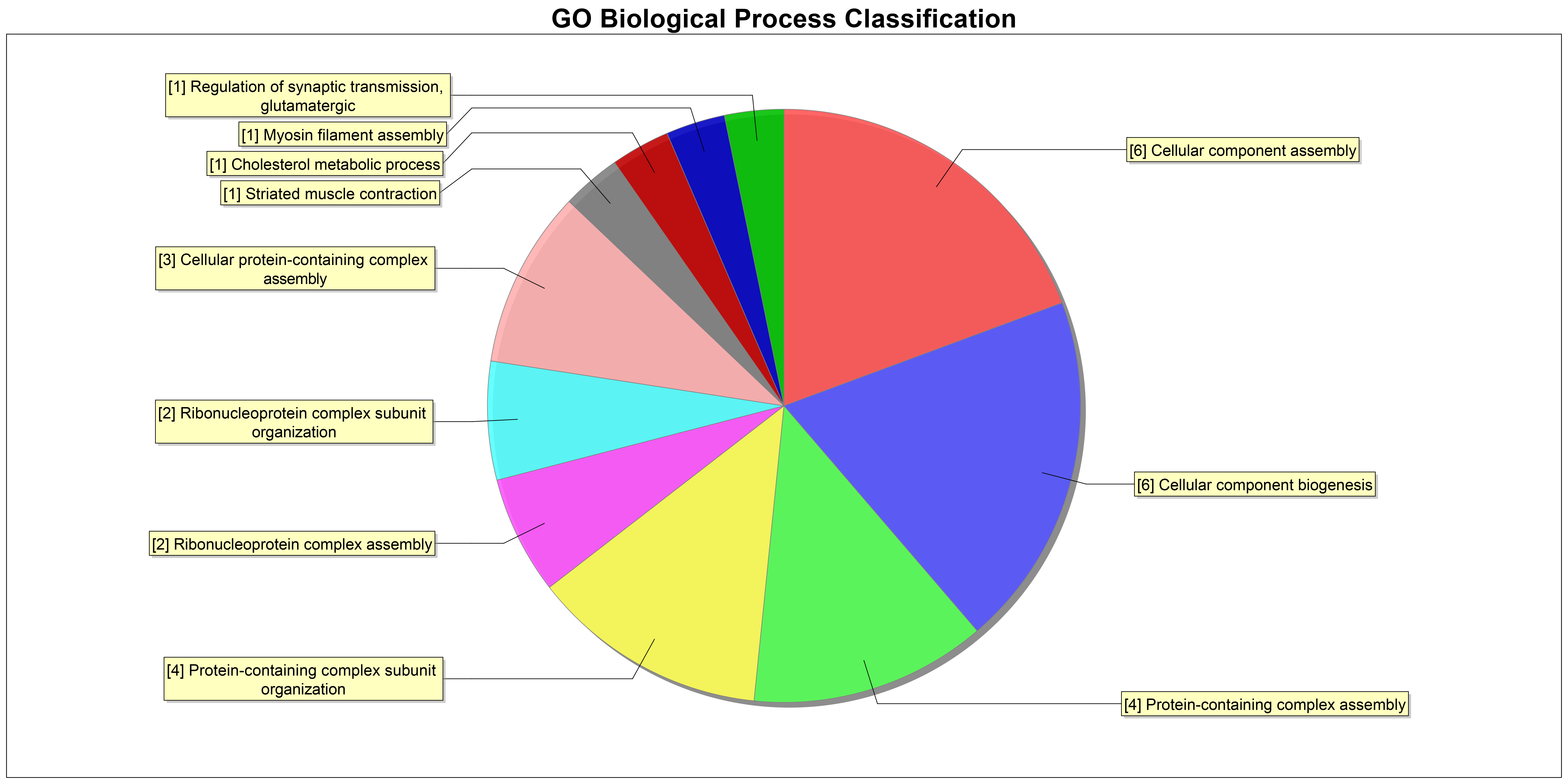

Supplementary Fig. 7 The KEGG analysis on differentially regulative protein

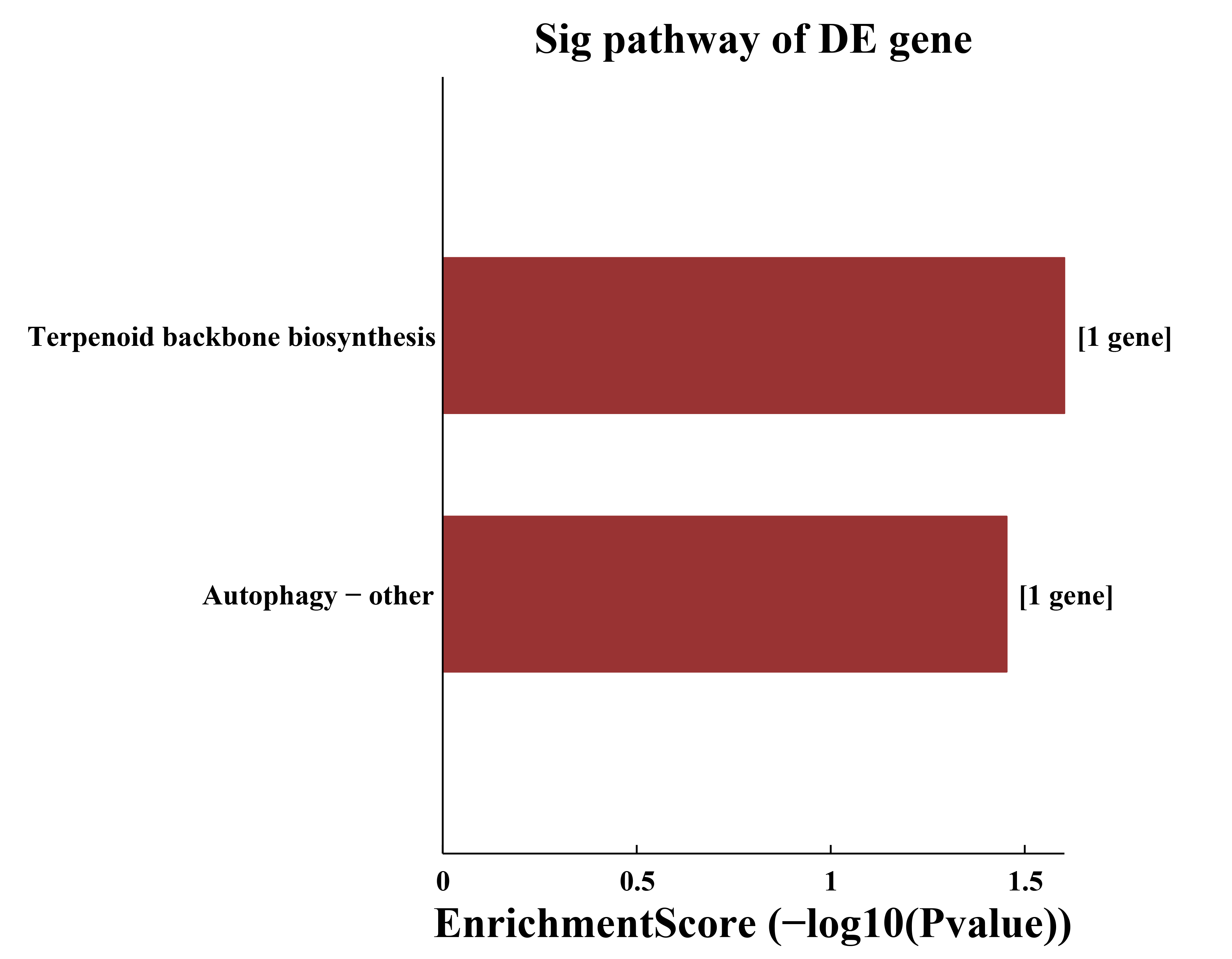

Supplementary Fig. 8 The PPI network according to the differentially regulative protein

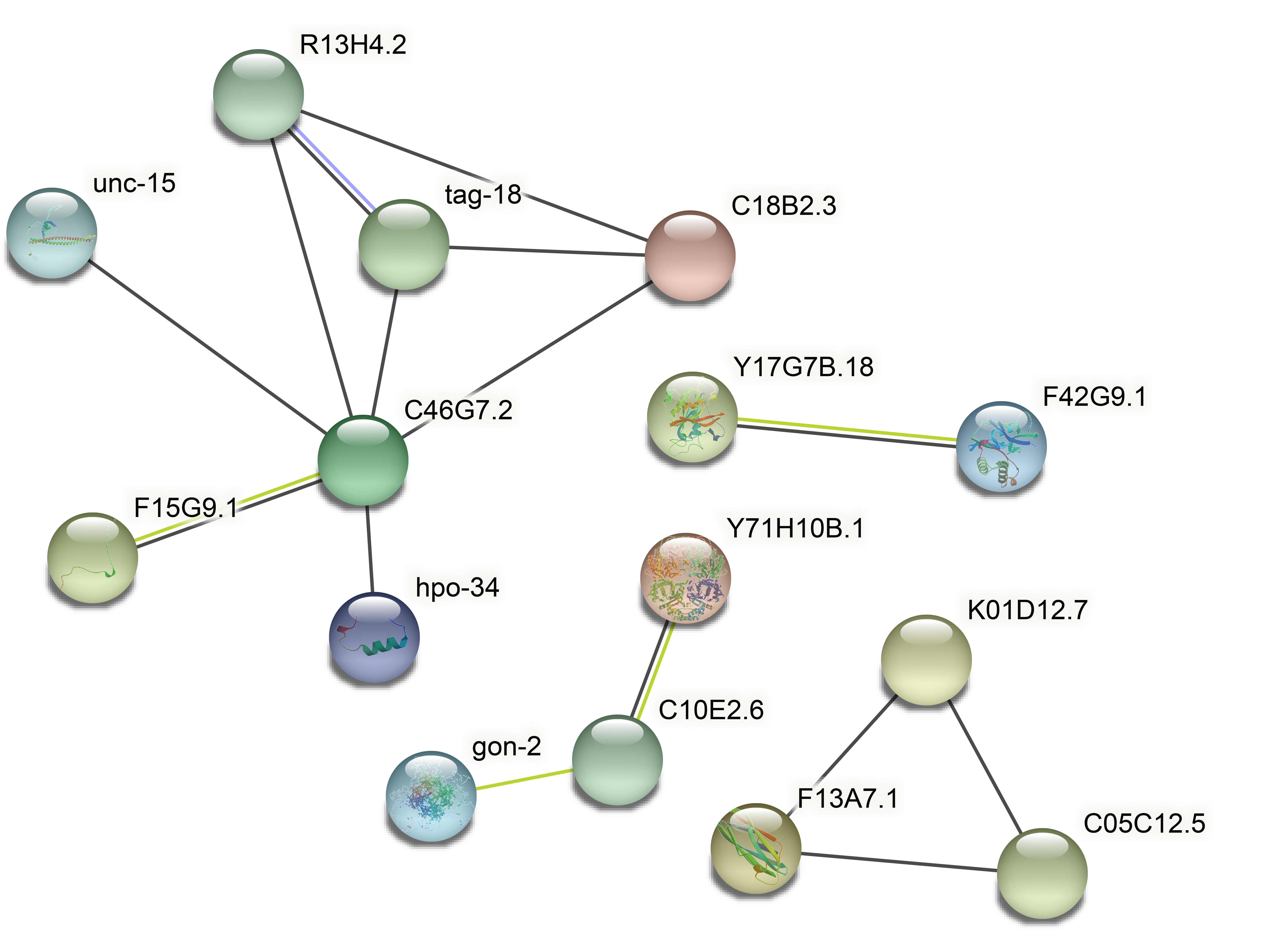

Supplementary Tabel 1 The prediction of miRNA-circRNA pairs.

| circRNA | direction | microRNA | direction |
| --- | --- | --- | --- |
| chrI:491196-494706:+ | up | cel-miR-1018 | down |
| cel_circ_0000130 | up | cel-miR-1819-3p | down |
| chrI:491196-494706:+ | up | cel-miR-1824-5p | up |
| chrI:491196-494706:+ | up | cel-miR-1830-3p | up |
| chrI:491196-494706:+ | up | cel-miR-1832a-5p | down |
| cel_circ_0000130 | up | cel-miR-2208b-5p | down |
| cel_circ_0000130 | up | cel-miR-2209a-3p | down |
| chrI:491196-494706:+ | up | cel-miR-2209a-3p | down |
| chrI:491196-494706:+ | up | cel-miR-2221 | up |
| chrI:491196-494706:+ | up | cel-miR-230-3p | up |
| cel_circ_0000130 | up | cel-miR-230-3p | up |
| chrI:491196-494706:+ | up | cel-miR-230-5p | up |
| chrI:491196-494706:+ | up | cel-miR-239a-5p | up |
| chrI:491196-494706:+ | up | cel-miR-243-3p | up |
| chrI:491196-494706:+ | up | cel-miR-243-5p | up |
| cel_circ_0000298 | up | cel-miR-243-5p | up |
| chrI:491196-494706:+ | up | cel-miR-249-3p | up |
| cel_circ_0000130 | up | cel-miR-253-3p | up |
| chrI:491196-494706:+ | up | cel-miR-35-3p | down |
| chrI:491196-494706:+ | up | cel-miR-36-3p | down |
| chrI:491196-494706:+ | up | cel-miR-37-3p | down |
| chrI:491196-494706:+ | up | cel-miR-39-3p | down |
| chrI:491196-494706:+ | up | cel-miR-39-5p | down |
| chrI:491196-494706:+ | up | cel-miR-40-3p | down |
| chrI:491196-494706:+ | up | cel-miR-41-3p | down |
| chrI:491196-494706:+ | up | cel-miR-41-5p | down |
| chrI:491196-494706:+ | up | cel-miR-42-3p | down |
| chrI:491196-494706:+ | up | cel-miR-45-5p | up |
| chrI:491196-494706:+ | up | cel-miR-4806-5p | down |
| chrI:491196-494706:+ | up | cel-miR-4807 | down |
| chrI:491196-494706:+ | up | cel-miR-4937 | up |
| cel_circ_0000130 | up | cel-miR-5546-3p | down |
| cel_circ_0000298 | up | cel-miR-5546-3p | down |
| chrI:491196-494706:+ | up | cel-miR-5546-3p | down |
| cel_circ_0000130 | up | cel-miR-58a-3p | up |
| chrI:491196-494706:+ | up | cel-miR-58a-3p | up |
| chrI:491196-494706:+ | up | cel-miR-58b-5p | down |
| chrI:491196-494706:+ | up | cel-miR-77-3p | down |
| cel_circ_0000130 | up | cel-miR-77-5p | down |
| chrI:491196-494706:+ | up | cel-miR-784-5p | down |
| chrI:491196-494706:+ | up | cel-miR-788-5p | up |
| cel_circ_0000130 | up | cel-miR-788-5p | up |
| chrI:491196-494706:+ | up | cel-miR-78 | down |
| chrI:491196-494706:+ | up | cel-miR-797-5p | up |
| chrI:491196-494706:+ | up | cel-miR-8200-3p | down |

Supplementary Table 2 The prediction of miRNA-lncRNA pairs.

| lncRNA | direction | microRNA | direction |
| --- | --- | --- | --- |
| K10G4.14 | up | cel-miR-2221 | up |
| K10G4.14 | up | cel-miR-237-5p | up |
| K10G4.14 | up | cel-miR-249-3p | up |
| C15H11.16 | up | cel-miR-4807 | down |
| K10G4.14 | up | cel-miR-4937 | up |
| C15H11.16 | up | cel-miR-5546-3p | down |
| K10G4.14 | up | cel-miR-58b-5p | down |
| K10G4.14 | up | cel-miR-784-5p | down |
| C15H11.16 | up | cel-miR-789-2-5p | down |
| K10G4.14 | up | cel-miR-797-5p | up |
| C15H11.16 | up | cel-miR-8196b-3p | down |

Supplementary Table 3 The prediction of miRNA-mRNA pairs of circRNA downstream.

| **microRNA** | **direction** | **mRNA** | **direction** | **parent gene** |
| --- | --- | --- | --- | --- |
| cel-miR-1018 | down | F58D5.5.1 | up | F58D5.5 |
| cel-miR-1018 | down | F40F8.1a.3 | up | F40F8.1 |
| cel-miR-1018 | down | C05D11.6a.1 | up | nas-4 |
| cel-miR-1018 | down | F53F10.2c.4 | up | F53F10.2 |
| cel-miR-1018 | down | F55A8.2g.1 | up | egl-4 |
| cel-miR-1018 | down | M88.6b.1 | up | pan-1 |
| cel-miR-1018 | down | M88.6a.1 | up | pan-1 |
| cel-miR-1018 | down | C04D8.1b.1 | up | pac-1 |
| cel-miR-1018 | down | R10E4.2q.1 | up | sup-26 |
| cel-miR-1018 | down | F38A6.3e.1 | up | hif-1 |
| cel-miR-1018 | down | Y67D8C.10d.3 | up | mca-3 |
| cel-miR-1018 | down | M03A1.6c.1 | up | ipla-1 |
| cel-miR-1018 | down | Y71G10AL.1a.1 | up | hrpu-2 |
| cel-miR-1018 | down | F09C3.2.1 | up | F09C3.2 |
| cel-miR-1018 | down | K10G6.3k.1 | up | sea-2 |
| cel-miR-1018 | down | C07D10.4.1 | up | nas-7 |
| cel-miR-1018 | down | Y75B8A.11a.1 | up | lury-1 |
| cel-miR-1018 | down | K10B2.1d.1 | up | lin-23 |
| cel-miR-1018 | down | F13B10.1b.1 | up | tir-1 |
| cel-miR-1018 | down | W01G7.1.1 | up | daf-5 |
| cel-miR-1018 | down | ZC250.1a.1 | up | cyn-17 |
| cel-miR-1018 | down | C48G7.3a.1 | up | rin-1 |
| cel-miR-1018 | down | F19H8.4.1 | up | mltn-9 |
| cel-miR-1018 | down | K01G5.9.3 | up | tasp-1 |
| cel-miR-1018 | down | C34G6.6a.1 | up | noah-1 |
| cel-miR-1018 | down | C34G6.6c.1 | up | noah-1 |
| cel-miR-1018 | down | F09F7.5a.1 | up | F09F7.5 |
| cel-miR-1018 | down | F49E11.1c.1 | up | mbk-2 |
| cel-miR-1018 | down | Y67D8C.5a.1 | up | eel-1 |
| cel-miR-1018 | down | C05D12.1.3 | up | C05D12.1 |
| cel-miR-1018 | down | C05D12.1.2 | up | C05D12.1 |
| cel-miR-1018 | down | F23B12.4a.2 | up | F23B12.4 |
| cel-miR-1018 | down | F26D12.1i.1 | up | fkh-7 |
| cel-miR-1018 | down | F26D12.1b.1 | up | fkh-7 |
| cel-miR-1018 | down | F26D12.1l.2 | up | fkh-7 |
| cel-miR-1018 | down | F26D12.1l.1 | up | fkh-7 |
| cel-miR-1018 | down | F26D12.1e.1 | up | fkh-7 |
| cel-miR-1018 | down | ZK783.1j.1 | up | fbn-1 |
| cel-miR-1018 | down | F56D2.1a.2 | up | ucr-1 |
| cel-miR-1018 | down | F56D2.1c.1 | up | ucr-1 |
| cel-miR-1018 | down | E01F3.1b.1 | up | pde-3 |
| cel-miR-1018 | down | R08F11.3.1 | up | cyp-33C8 |
| cel-miR-1018 | down | R09B5.9.1 | up | cnc-4 |
| cel-miR-1018 | down | K09E2.4b.3 | up | rig-1 |
| cel-miR-1018 | down | C07G1.1.1 | up | svh-1 |
| cel-miR-1018 | down | D2023.1f.1 | up | D2023.1 |
| cel-miR-1018 | down | R13H7.2d.1 | up | R13H7.2 |
| cel-miR-1018 | down | F19B10.1.2 | up | F19B10.1 |
| cel-miR-1819-3p | down | C01B7.4a.1 | up | magu-2 |
| cel-miR-1819-3p | down | C01B7.4a.2 | up | magu-2 |
| cel-miR-1819-3p | down | C24A3.2a.1 | up | C24A3.2 |
| cel-miR-1819-3p | down | H14N18.1c.2 | up | unc-23 |
| cel-miR-1819-3p | down | H14N18.1b.1 | up | unc-23 |
| cel-miR-1819-3p | down | H14N18.1b.3 | up | unc-23 |
| cel-miR-1819-3p | down | H14N18.1a.1 | up | unc-23 |
| cel-miR-1819-3p | down | H14N18.1b.2 | up | unc-23 |
| cel-miR-1819-3p | down | F54C9.1.2 | up | iff-2 |
| cel-miR-1819-3p | down | ZK742.1a.3 | up | xpo-1 |
| cel-miR-1819-3p | down | C34G6.6a.1 | up | noah-1 |
| cel-miR-1819-3p | down | C34G6.6c.1 | up | noah-1 |
| cel-miR-1819-3p | down | ZK1290.8.1 | up | wrt-10 |
| cel-miR-1819-3p | down | C56E6.2.1 | up | C56E6.2 |
| cel-miR-1819-3p | down | F35C8.6.2 | up | pfn-2 |
| cel-miR-1819-3p | down | Y76B12C.9.1 | up | Y76B12C.9 |
| cel-miR-1819-3p | down | B0491.8b.1 | up | clh-2 |
| cel-miR-1819-3p | down | C02E11.1a.1 | up | nra-4 |
| cel-miR-1819-3p | down | K09A9.2.1 | up | rab-14 |
| cel-miR-1819-3p | down | W02C12.3b.1 | up | hlh-30 |
| cel-miR-1819-3p | down | R11.2.1 | up | nlp-67 |
| cel-miR-1819-3p | down | C33A12.3a.1 | up | C33A12.3 |
| cel-miR-1819-3p | down | Y39E4B.12a.2 | up | gly-5 |
| cel-miR-1819-3p | down | Y39E4B.12c.2 | up | gly-5 |
| cel-miR-1819-3p | down | F18A1.2a.1 | up | lin-26 |
| cel-miR-1819-3p | down | F53F1.4.1 | up | F53F1.4 |
| cel-miR-1819-3p | down | F42C5.7.1 | up | grl-4 |
| cel-miR-1819-3p | down | C16D6.2a.1 | up | npr-4 |
| cel-miR-1819-3p | down | F13C5.1.1 | up | F13C5.1 |
| cel-miR-1819-3p | down | C14B9.4b.1 | up | plk-1 |
| cel-miR-1819-3p | down | F53G12.5b.1 | up | mex-3 |
| cel-miR-1819-3p | down | C38D4.6c.4 | up | pal-1 |
| cel-miR-1819-3p | down | C05D12.7.1 | up | C05D12.7 |
| cel-miR-1819-3p | down | W04G3.3.1 | up | lpr-4 |
| cel-miR-1819-3p | down | F55F3.3b.1 | up | nkb-3 |
| cel-miR-1819-3p | down | W06F12.1b.2 | up | lit-1 |
| cel-miR-1819-3p | down | W06F12.1c.1 | up | lit-1 |
| cel-miR-1819-3p | down | C42D4.18.1 | up | C42D4.18 |
| cel-miR-1819-3p | down | F22D3.1b.1 | up | ceh-38 |
| cel-miR-1819-3p | down | F22D3.1c.2 | up | ceh-38 |
| cel-miR-1819-3p | down | F22D3.1d.2 | up | ceh-38 |
| cel-miR-1819-3p | down | F22D3.1e.1 | up | ceh-38 |
| cel-miR-1819-3p | down | F22D3.1f.1 | up | ceh-38 |
| cel-miR-1819-3p | down | F22D3.1g.1 | up | ceh-38 |
| cel-miR-1819-3p | down | F22D3.1h.1 | up | ceh-38 |
| cel-miR-1819-3p | down | F22D3.1i.1 | up | ceh-38 |
| cel-miR-1819-3p | down | F22D3.1j.1 | up | ceh-38 |
| cel-miR-1819-3p | down | F22D3.1k.1 | up | ceh-38 |
| cel-miR-1819-3p | down | F22D3.1m.1 | up | ceh-38 |
| cel-miR-1819-3p | down | F22D3.1n.1 | up | ceh-38 |
| cel-miR-1819-3p | down | F22D3.1p.1 | up | ceh-38 |
| cel-miR-1819-3p | down | F46G10.6.1 | up | mxl-3 |
| cel-miR-1819-3p | down | R07E3.2.1 | up | R07E3.2 |
| cel-miR-1819-3p | down | K05B2.2b.1 | up | K05B2.2 |
| cel-miR-1819-3p | down | F53A10.2d.1 | up | F53A10.2 |
| cel-miR-1819-3p | down | C07A12.1b.1 | up | ham-2 |
| cel-miR-1819-3p | down | C38H2.2.2 | up | C38H2.2 |
| cel-miR-1819-3p | down | ZC518.3d.1 | up | ccr-4 |
| cel-miR-1819-3p | down | Y73B6A.5c.1 | up | lin-45 |
| cel-miR-1819-3p | down | F52E10.4c.1 | up | F52E10.4 |
| cel-miR-1819-3p | down | C05C8.7.4 | up | C05C8.7 |
| cel-miR-1819-3p | down | C05C8.7.1 | up | C05C8.7 |
| cel-miR-1819-3p | down | M60.7.1 | up | M60.7 |
| cel-miR-1819-3p | down | ZK54.2b.1 | up | tps-1 |
| cel-miR-1819-3p | down | Y38C1BA.3.1 | up | col-109 |
| cel-miR-1819-3p | down | F37C4.4b.2 | up | F37C4.4 |
| cel-miR-1819-3p | down | F37C4.4b.1 | up | F37C4.4 |
| cel-miR-1819-3p | down | C15F1.2.1 | up | C15F1.2 |
| cel-miR-1819-3p | down | Y81G3A.3a.2 | up | gcn-2 |
| cel-miR-1819-3p | down | D1054.9e.1 | up | D1054.9 |
| cel-miR-1819-3p | down | D1054.9d.1 | up | D1054.9 |
| cel-miR-1819-3p | down | D1054.9c.1 | up | D1054.9 |
| cel-miR-1819-3p | down | D1054.9b.1 | up | D1054.9 |
| cel-miR-1819-3p | down | D1054.9a.1 | up | D1054.9 |
| cel-miR-1819-3p | down | F35H10.4.3 | up | vha-5 |
| cel-miR-1819-3p | down | F35H10.4.1 | up | vha-5 |
| cel-miR-1819-3p | down | ZK270.2b.1 | up | frm-1 |
| cel-miR-1819-3p | down | Y92C3B.2d.1 | up | uaf-1 |
| cel-miR-1819-3p | down | Y92C3B.2c.1 | up | uaf-1 |
| cel-miR-1819-3p | down | D1005.3.2 | up | cebp-1 |
| cel-miR-1819-3p | down | D1005.3.1 | up | cebp-1 |
| cel-miR-1819-3p | down | F59B10.1.1 | up | myrf-1 |
| cel-miR-1819-3p | down | C17C3.12c.1 | up | acdh-2 |
| cel-miR-1819-3p | down | F01F1.8b.2 | up | cct-6 |
| cel-miR-1819-3p | down | F01F1.8b.1 | up | cct-6 |
| cel-miR-1819-3p | down | ZK1005.1a.1 | up | tank-1 |
| cel-miR-1819-3p | down | ZK970.1b.1 | up | nep-26 |
| cel-miR-1819-3p | down | C05D10.4b.1 | up | C05D10.4 |
| cel-miR-1819-3p | down | K08B12.2a.1 | up | dmd-7 |
| cel-miR-1819-3p | down | C37F5.1b.1 | up | lin-1 |
| cel-miR-1819-3p | down | Y64G10A.7c.1 | up | Y64G10A.7 |
| cel-miR-1819-3p | down | F37A8.5.1 | up | F37A8.5 |
| cel-miR-1819-3p | down | Y73B6BL.31b.1 | up | Y73B6BL.31 |
| cel-miR-1819-3p | down | M02A10.3b.1 | up | sli-1 |
| cel-miR-1819-3p | down | C04A2.3p.1 | up | egl-27 |
| cel-miR-1819-3p | down | F56H1.1.1 | up | che-14 |
| cel-miR-1819-3p | down | F31F7.1b.2 | up | F31F7.1 |
| cel-miR-1819-3p | down | F43C9.2b.1 | up | F43C9.2 |
| cel-miR-1819-3p | down | Y38E10A.15.1 | up | nspe-7 |
| cel-miR-1819-3p | down | K07C5.9.1 | up | K07C5.9 |
| cel-miR-1819-3p | down | F55A11.4a.2 | up | F55A11.4 |
| cel-miR-1819-3p | down | C12D8.11.1 | up | rop-1 |
| cel-miR-1819-3p | down | C12D8.11.2 | up | rop-1 |
| cel-miR-1819-3p | down | C34H3.2.1 | up | odd-2 |
| cel-miR-1819-3p | down | Y26E6A.2.1 | up | Y26E6A.2 |
| cel-miR-1819-3p | down | H32C10.2b.1 | up | lin-33 |
| cel-miR-1819-3p | down | F22E5.13.1 | up | F22E5.13 |
| cel-miR-1819-3p | down | F22E5.13.2 | up | F22E5.13 |
| cel-miR-1819-3p | down | ZC239.6.3 | up | ZC239.6 |
| cel-miR-1819-3p | down | ZC239.6.1 | up | ZC239.6 |
| cel-miR-1819-3p | down | ZC239.6.2 | up | ZC239.6 |
| cel-miR-1819-3p | down | F29C12.1a.1 | up | pqn-32 |
| cel-miR-1819-3p | down | F41G4.7.1 | up | F41G4.7 |
| cel-miR-1819-3p | down | Y41E3.2.1 | up | dpy-4 |
| cel-miR-1819-3p | down | W01G7.1.1 | up | daf-5 |
| cel-miR-1819-3p | down | K03H1.10.5 | up | cbp-2 |
| cel-miR-1819-3p | down | F40F12.7.2 | up | cbp-3 |
| cel-miR-1819-3p | down | ZK287.2a.3 | up | sulp-8 |
| cel-miR-1819-3p | down | Y38H8A.1.1 | up | Y38H8A.1 |
| cel-miR-1819-3p | down | C23H4.6b.1 | up | C23H4.6 |
| cel-miR-1819-3p | down | C23H4.6a.1 | up | C23H4.6 |
| cel-miR-1819-3p | down | C02F5.8.1 | up | tsp-1 |
| cel-miR-1819-3p | down | F46C8.6.1 | up | dpy-7 |
| cel-miR-1819-3p | down | F40E10.5.1 | up | F40E10.5 |
| cel-miR-1832a-5p | down | Y111B2A.18.1 | up | rsp-3 |
| cel-miR-2208b-5p | down | F58D5.5.1 | up | F58D5.5 |
| cel-miR-2208b-5p | down | F29C4.8d.1 | up | col-99 |
| cel-miR-2208b-5p | down | H11L12.1.1 | up | H11L12.1 |
| cel-miR-2208b-5p | down | Y54G2A.29.1 | up | ndnf-1 |
| cel-miR-2208b-5p | down | D2023.1f.1 | up | D2023.1 |
| cel-miR-2208b-5p | down | C02F12.1b.1 | up | tsp-17 |
| cel-miR-2209a-3p | down | F35G8.1.1 | up | npr-7 |
| cel-miR-2209a-3p | down | C41D11.1b.1 | up | eri-6 |
| cel-miR-2209a-3p | down | F46G10.2.1 | up | F46G10.2 |
| cel-miR-2209a-3p | down | F52E1.13b.2 | up | lmd-3 |
| cel-miR-2209a-3p | down | Y22D7AL.8d.1 | up | sms-3 |
| cel-miR-2209a-3p | down | F37A8.5.1 | up | F37A8.5 |
| cel-miR-2209a-3p | down | M03F4.7a.2 | up | calu-1 |
| cel-miR-2209a-3p | down | H19M22.3a.1 | up | zmp-2 |
| cel-miR-2209a-3p | down | H19M22.3c.1 | up | zmp-2 |
| cel-miR-2209a-3p | down | F28B12.1b.1 | up | F28B12.1 |
| cel-miR-2209a-3p | down | F28B12.1d.1 | up | F28B12.1 |
| cel-miR-2209a-3p | down | F28B12.1d.2 | up | F28B12.1 |
| cel-miR-2209a-3p | down | K02E10.2b.1 | up | hid-1 |
| cel-miR-2209a-3p | down | B0213.4.1 | up | nlp-29 |
| cel-miR-2209a-3p | down | F13B10.1b.1 | up | tir-1 |
| cel-miR-2209a-3p | down | F22E12.4d.2 | up | egl-9 |
| cel-miR-2209a-3p | down | Y37A1B.7.1 | up | Y37A1B.7 |
| cel-miR-2209a-3p | down | F09B9.1a.1 | up | oac-14 |
| cel-miR-2209a-3p | down | M03F8.1.1 | up | M03F8.1 |
| cel-miR-2209a-3p | down | K10G6.3k.1 | up | sea-2 |
| cel-miR-2209a-3p | down | F52F12.1b.1 | up | oct-1 |
| cel-miR-2209a-3p | down | C05D11.6a.1 | up | nas-4 |
| cel-miR-2209a-3p | down | Y38E10A.15.1 | up | nspe-7 |
| cel-miR-2209a-3p | down | F28F5.6.1 | up | F28F5.6 |
| cel-miR-2209a-3p | down | ZK20.1.1 | up | ghi-1 |
| cel-miR-2209a-3p | down | C07D10.4.1 | up | nas-7 |
| cel-miR-2209a-3p | down | R11A5.7.1 | up | suro-1 |
| cel-miR-2209a-3p | down | ZK180.5a.1 | up | ZK180.5 |
| cel-miR-2209a-3p | down | ZK180.5b.1 | up | ZK180.5 |
| cel-miR-2209a-3p | down | C54G10.4b.1 | up | slc-25A29 |
| cel-miR-2209a-3p | down | F01D4.6a.1 | up | mec-3 |
| cel-miR-2209a-3p | down | K08B12.2a.1 | up | dmd-7 |
| cel-miR-2209a-3p | down | F58F9.1a.1 | up | F58F9.1 |
| cel-miR-2209a-3p | down | C09F12.3.1 | up | C09F12.3 |
| cel-miR-2209a-3p | down | F13B12.5.1 | up | ins-1 |
| cel-miR-2209a-3p | down | ZC250.4.1 | up | ZC250.4 |
| cel-miR-2209a-3p | down | Y39B6A.19a.1 | up | twk-46 |
| cel-miR-2209a-3p | down | H23N18.3.1 | up | ugt-8 |
| cel-miR-2209a-3p | down | C01F1.5.1 | up | C01F1.5 |
| cel-miR-2209a-3p | down | C40C9.5b.1 | up | nlg-1 |
| cel-miR-2209a-3p | down | C06E2.1.1 | up | C06E2.1 |
| cel-miR-2209a-3p | down | F11E6.9.1 | up | F11E6.9 |
| cel-miR-2209a-3p | down | C05E11.7.1 | up | C05E11.7 |
| cel-miR-2209a-3p | down | ZK75.1.1 | up | ins-4 |
| cel-miR-35-3p | down | ZK637.8b.1 | up | unc-32 |
| cel-miR-35-3p | down | C34F11.3c.1 | up | ampd-1 |
| cel-miR-35-3p | down | Y67D8C.10d.3 | up | mca-3 |
| cel-miR-35-3p | down | Y55F3AM.10.1 | up | abhd-14 |
| cel-miR-35-3p | down | F35H10.4.3 | up | vha-5 |
| cel-miR-35-3p | down | F35H10.4.1 | up | vha-5 |
| cel-miR-35-3p | down | Y43F4A.1a.1 | up | Y43F4A.1 |
| cel-miR-35-3p | down | ZK1321.2a.1 | up | shk-1 |
| cel-miR-35-3p | down | W03C9.4c.2 | up | lin-29 |
| cel-miR-35-3p | down | W03C9.4c.1 | up | lin-29 |
| cel-miR-35-3p | down | W03C9.4a.1 | up | lin-29 |
| cel-miR-35-3p | down | C48D5.3.1 | up | C48D5.3 |
| cel-miR-35-3p | down | M03F8.1.1 | up | M03F8.1 |
| cel-miR-35-3p | down | K04G11.5c.1 | up | irk-3 |
| cel-miR-35-3p | down | K04G11.5e.1 | up | irk-3 |
| cel-miR-35-3p | down | ZK455.7.2 | up | pgp-3 |
| cel-miR-35-3p | down | F27D9.7.1 | up | F27D9.7 |
| cel-miR-35-3p | down | R03H4.6.1 | up | bus-1 |
| cel-miR-35-3p | down | Y67A10A.8a.1 | up | paqr-3 |
| cel-miR-35-3p | down | C34F11.9f.1 | up | dsh-1 |
| cel-miR-35-3p | down | F32D8.1.1 | up | F32D8.1 |
| cel-miR-35-3p | down | F29B9.8.2 | up | F29B9.8 |
| cel-miR-35-3p | down | H10E21.5.1 | up | H10E21.5 |
| cel-miR-35-3p | down | F58A4.7b.2 | up | hlh-11 |
| cel-miR-36-3p | down | ZK637.8b.1 | up | unc-32 |
| cel-miR-36-3p | down | C34F11.3c.1 | up | ampd-1 |
| cel-miR-36-3p | down | Y67D8C.10d.3 | up | mca-3 |
| cel-miR-36-3p | down | Y55F3AM.10.1 | up | abhd-14 |
| cel-miR-36-3p | down | F35H10.4.3 | up | vha-5 |
| cel-miR-36-3p | down | F35H10.4.1 | up | vha-5 |
| cel-miR-36-3p | down | ZK1321.2a.1 | up | shk-1 |
| cel-miR-36-3p | down | W03C9.4c.2 | up | lin-29 |
| cel-miR-36-3p | down | W03C9.4c.1 | up | lin-29 |
| cel-miR-36-3p | down | W03C9.4a.1 | up | lin-29 |
| cel-miR-36-3p | down | M03F8.1.1 | up | M03F8.1 |
| cel-miR-36-3p | down | K04G11.5c.1 | up | irk-3 |
| cel-miR-36-3p | down | K04G11.5e.1 | up | irk-3 |
| cel-miR-36-3p | down | ZK455.7.2 | up | pgp-3 |
| cel-miR-36-3p | down | F27D9.7.1 | up | F27D9.7 |
| cel-miR-36-3p | down | R03H4.6.1 | up | bus-1 |
| cel-miR-36-3p | down | Y67A10A.8a.1 | up | paqr-3 |
| cel-miR-36-3p | down | C34F11.9f.1 | up | dsh-1 |
| cel-miR-36-3p | down | F32D8.1.1 | up | F32D8.1 |
| cel-miR-36-3p | down | F29B9.8.2 | up | F29B9.8 |
| cel-miR-36-3p | down | H10E21.5.1 | up | H10E21.5 |
| cel-miR-36-3p | down | F58A4.7b.2 | up | hlh-11 |
| cel-miR-37-3p | down | C34F11.3c.1 | up | ampd-1 |
| cel-miR-37-3p | down | Y67D8C.10d.3 | up | mca-3 |
| cel-miR-37-3p | down | Y55F3AM.10.1 | up | abhd-14 |
| cel-miR-37-3p | down | F35H10.4.3 | up | vha-5 |
| cel-miR-37-3p | down | F35H10.4.1 | up | vha-5 |
| cel-miR-37-3p | down | ZK1321.2a.1 | up | shk-1 |
| cel-miR-37-3p | down | W03C9.4c.2 | up | lin-29 |
| cel-miR-37-3p | down | W03C9.4c.1 | up | lin-29 |
| cel-miR-37-3p | down | W03C9.4a.1 | up | lin-29 |
| cel-miR-37-3p | down | M03F8.1.1 | up | M03F8.1 |
| cel-miR-37-3p | down | K04G11.5c.1 | up | irk-3 |
| cel-miR-37-3p | down | K04G11.5e.1 | up | irk-3 |
| cel-miR-37-3p | down | ZK455.7.2 | up | pgp-3 |
| cel-miR-37-3p | down | F27D9.7.1 | up | F27D9.7 |
| cel-miR-37-3p | down | R03H4.6.1 | up | bus-1 |
| cel-miR-37-3p | down | Y67A10A.8a.1 | up | paqr-3 |
| cel-miR-37-3p | down | C34F11.9f.1 | up | dsh-1 |
| cel-miR-37-3p | down | F32D8.1.1 | up | F32D8.1 |
| cel-miR-37-3p | down | F29B9.8.2 | up | F29B9.8 |
| cel-miR-37-3p | down | F58A4.7b.2 | up | hlh-11 |
| cel-miR-39-3p | down | C34F11.3c.1 | up | ampd-1 |
| cel-miR-39-3p | down | Y55F3AM.10.1 | up | abhd-14 |
| cel-miR-39-3p | down | F35H10.4.3 | up | vha-5 |
| cel-miR-39-3p | down | F35H10.4.1 | up | vha-5 |
| cel-miR-39-3p | down | ZK1321.2a.1 | up | shk-1 |
| cel-miR-39-3p | down | W03C9.4c.2 | up | lin-29 |
| cel-miR-39-3p | down | W03C9.4c.1 | up | lin-29 |
| cel-miR-39-3p | down | W03C9.4a.1 | up | lin-29 |
| cel-miR-39-3p | down | M03F8.1.1 | up | M03F8.1 |
| cel-miR-39-3p | down | K04G11.5c.1 | up | irk-3 |
| cel-miR-39-3p | down | K04G11.5e.1 | up | irk-3 |
| cel-miR-39-3p | down | ZK455.7.2 | up | pgp-3 |
| cel-miR-39-3p | down | R03H4.6.1 | up | bus-1 |
| cel-miR-39-3p | down | Y67A10A.8a.1 | up | paqr-3 |
| cel-miR-39-3p | down | C34F11.9f.1 | up | dsh-1 |
| cel-miR-39-3p | down | F32D8.1.1 | up | F32D8.1 |
| cel-miR-39-3p | down | F29B9.8.2 | up | F29B9.8 |
| cel-miR-39-3p | down | H10E21.5.1 | up | H10E21.5 |
| cel-miR-39-3p | down | F58A4.7b.2 | up | hlh-11 |
| cel-miR-39-5p | down | K11H12.3.1 | up | K11H12.3 |
| cel-miR-39-5p | down | Y50E8A.16.1 | up | haf-7 |
| cel-miR-39-5p | down | R06C7.5b.1 | up | adsl-1 |
| cel-miR-40-3p | down | C34F11.3c.1 | up | ampd-1 |
| cel-miR-40-3p | down | Y67D8C.10d.3 | up | mca-3 |
| cel-miR-40-3p | down | Y55F3AM.10.1 | up | abhd-14 |
| cel-miR-40-3p | down | ZK1321.2a.1 | up | shk-1 |
| cel-miR-40-3p | down | W03C9.4c.2 | up | lin-29 |
| cel-miR-40-3p | down | W03C9.4c.1 | up | lin-29 |
| cel-miR-40-3p | down | W03C9.4a.1 | up | lin-29 |
| cel-miR-40-3p | down | M03F8.1.1 | up | M03F8.1 |
| cel-miR-40-3p | down | K04G11.5c.1 | up | irk-3 |
| cel-miR-40-3p | down | K04G11.5e.1 | up | irk-3 |
| cel-miR-40-3p | down | ZK455.7.2 | up | pgp-3 |
| cel-miR-40-3p | down | Y67A10A.8a.1 | up | paqr-3 |
| cel-miR-40-3p | down | C34F11.9f.1 | up | dsh-1 |
| cel-miR-40-3p | down | F32D8.1.1 | up | F32D8.1 |
| cel-miR-40-3p | down | F29B9.8.2 | up | F29B9.8 |
| cel-miR-40-3p | down | F58A4.7b.2 | up | hlh-11 |
| cel-miR-41-3p | down | Y55F3AM.10.1 | up | abhd-14 |
| cel-miR-41-3p | down | F35H10.4.3 | up | vha-5 |
| cel-miR-41-3p | down | F35H10.4.1 | up | vha-5 |
| cel-miR-41-3p | down | ZK1321.2a.1 | up | shk-1 |
| cel-miR-41-3p | down | W03C9.4c.2 | up | lin-29 |
| cel-miR-41-3p | down | W03C9.4c.1 | up | lin-29 |
| cel-miR-41-3p | down | W03C9.4a.1 | up | lin-29 |
| cel-miR-41-3p | down | M03F8.1.1 | up | M03F8.1 |
| cel-miR-41-3p | down | K04G11.5c.1 | up | irk-3 |
| cel-miR-41-3p | down | K04G11.5e.1 | up | irk-3 |
| cel-miR-41-3p | down | ZK455.7.2 | up | pgp-3 |
| cel-miR-41-3p | down | F27D9.7.1 | up | F27D9.7 |
| cel-miR-41-3p | down | R03H4.6.1 | up | bus-1 |
| cel-miR-41-3p | down | Y67A10A.8a.1 | up | paqr-3 |
| cel-miR-41-3p | down | C34F11.9f.1 | up | dsh-1 |
| cel-miR-41-3p | down | F32D8.1.1 | up | F32D8.1 |
| cel-miR-41-3p | down | F29B9.8.2 | up | F29B9.8 |
| cel-miR-41-3p | down | H10E21.5.1 | up | H10E21.5 |
| cel-miR-41-3p | down | F58A4.7b.2 | up | hlh-11 |
| cel-miR-41-5p | down | Y47G6A.19b.1 | up | Y47G6A.19 |
| cel-miR-41-5p | down | C37A2.5b.2 | up | pqn-21 |
| cel-miR-41-5p | down | B0412.2.1 | up | daf-7 |
| cel-miR-41-5p | down | ZC239.6.3 | up | ZC239.6 |
| cel-miR-41-5p | down | H18N23.2c.1 | up | H18N23.2 |
| cel-miR-41-5p | down | C10C5.1k.1 | up | pezo-1 |
| cel-miR-41-5p | down | C10C5.1a.1 | up | pezo-1 |
| cel-miR-41-5p | down | F47F6.1b.1 | up | lin-42 |
| cel-miR-41-5p | down | F31D4.8.1 | up | F31D4.8 |
| cel-miR-41-5p | down | M163.1.1 | up | M163.1 |
| cel-miR-41-5p | down | F55F3.3b.1 | up | nkb-3 |
| cel-miR-41-5p | down | R153.1h.1 | up | pde-4 |
| cel-miR-41-5p | down | R153.1a.1 | up | pde-4 |
| cel-miR-41-5p | down | C28G1.5a.1 | up | C28G1.5 |
| cel-miR-41-5p | down | F21G4.5.1 | up | F21G4.5 |
| cel-miR-41-5p | down | F41H10.11.1 | up | sand-1 |
| cel-miR-41-5p | down | M28.10.1 | up | M28.10 |
| cel-miR-41-5p | down | K02E10.2b.1 | up | hid-1 |
| cel-miR-41-5p | down | R02F2.1a.3 | up | R02F2.1 |
| cel-miR-41-5p | down | R02F2.1b.3 | up | R02F2.1 |
| cel-miR-41-5p | down | F08B12.1.1 | up | prmn-1 |
| cel-miR-41-5p | down | Y65B4BM.3.1 | up | Y65B4BM.3 |
| cel-miR-41-5p | down | F40F9.2.2 | up | tag-120 |
| cel-miR-41-5p | down | B0495.7.3 | up | B0495.7 |
| cel-miR-41-5p | down | F01G12.5b.1 | up | let-2 |
| cel-miR-41-5p | down | F59B10.1.1 | up | myrf-1 |
| cel-miR-41-5p | down | B0280.4.1 | up | odd-1 |
| cel-miR-41-5p | down | Y45F10B.13a.1 | up | Y45F10B.13 |
| cel-miR-41-5p | down | Y45F10B.13b.1 | up | Y45F10B.13 |
| cel-miR-41-5p | down | F29C12.1a.1 | up | pqn-32 |
| cel-miR-41-5p | down | ZC250.1a.1 | up | cyn-17 |
| cel-miR-41-5p | down | F09F7.5a.1 | up | F09F7.5 |
| cel-miR-41-5p | down | F39D8.1b.1 | up | pqn-36 |
| cel-miR-41-5p | down | R90.3.1 | up | ttr-28 |
| cel-miR-41-5p | down | F14D2.19.1 | up | F14D2.19 |
| cel-miR-42-3p | down | C34F11.3c.1 | up | ampd-1 |
| cel-miR-42-3p | down | Y55F3AM.10.1 | up | abhd-14 |
| cel-miR-42-3p | down | F35H10.4.3 | up | vha-5 |
| cel-miR-42-3p | down | F35H10.4.1 | up | vha-5 |
| cel-miR-42-3p | down | ZK1321.2a.1 | up | shk-1 |
| cel-miR-42-3p | down | W03C9.4c.2 | up | lin-29 |
| cel-miR-42-3p | down | W03C9.4c.1 | up | lin-29 |
| cel-miR-42-3p | down | W03C9.4a.1 | up | lin-29 |
| cel-miR-42-3p | down | M03F8.1.1 | up | M03F8.1 |
| cel-miR-42-3p | down | ZK455.7.2 | up | pgp-3 |
| cel-miR-42-3p | down | F27D9.7.1 | up | F27D9.7 |
| cel-miR-42-3p | down | R03H4.6.1 | up | bus-1 |
| cel-miR-42-3p | down | Y67A10A.8a.1 | up | paqr-3 |
| cel-miR-42-3p | down | C34F11.9f.1 | up | dsh-1 |
| cel-miR-42-3p | down | F32D8.1.1 | up | F32D8.1 |
| cel-miR-42-3p | down | F29B9.8.2 | up | F29B9.8 |
| cel-miR-42-3p | down | F58A4.7b.2 | up | hlh-11 |
| cel-miR-4806-5p | down | F35C8.6.2 | up | pfn-2 |
| cel-miR-4807 | down | R03G5.1c.1 | up | eef-1A.2 |
| cel-miR-4807 | down | F58D5.5.1 | up | F58D5.5 |
| cel-miR-4807 | down | F21A10.2d.1 | up | myrf-2 |
| cel-miR-4807 | down | K11E8.1d.1 | up | unc-43 |
| cel-miR-4807 | down | K11E8.1e.1 | up | unc-43 |
| cel-miR-4807 | down | K11E8.1o.1 | up | unc-43 |
| cel-miR-4807 | down | K11E8.1r.1 | up | unc-43 |
| cel-miR-4807 | down | C25E10.5.1 | up | C25E10.5 |
| cel-miR-4807 | down | Y37A1B.2h.3 | up | lst-4 |
| cel-miR-4807 | down | Y37E3.16.2 | up | fpn-1.1 |
| cel-miR-4807 | down | Y37E3.16.1 | up | fpn-1.1 |
| cel-miR-4807 | down | C44C10.11.1 | up | C44C10.11 |
| cel-miR-4807 | down | F49E11.1c.1 | up | mbk-2 |
| cel-miR-4807 | down | ZK177.8b.2 | up | ZK177.8 |
| cel-miR-4807 | down | C46C2.1n.2 | up | wnk-1 |
| cel-miR-4807 | down | C46C2.1d.1 | up | wnk-1 |
| cel-miR-4807 | down | F30A10.2.1 | up | F30A10.2 |
| cel-miR-4807 | down | F55H12.4.1 | up | F55H12.4 |
| cel-miR-4807 | down | R03G5.1a.3 | up | eef-1A.2 |
| cel-miR-4807 | down | R03G5.1a.2 | up | eef-1A.2 |
| cel-miR-4807 | down | R03G5.1d.2 | up | eef-1A.2 |
| cel-miR-4807 | down | R03G5.1d.1 | up | eef-1A.2 |
| cel-miR-4807 | down | R03G5.1b.1 | up | eef-1A.2 |
| cel-miR-4807 | down | F55A11.4a.2 | up | F55A11.4 |
| cel-miR-5546-3p | down | R166.1a.1 | up | mab-10 |
| cel-miR-5546-3p | down | Y55B1BL.1a.1 | up | Y55B1BL.1 |
| cel-miR-5546-3p | down | K04G11.5c.1 | up | irk-3 |
| cel-miR-5546-3p | down | K04G11.5e.1 | up | irk-3 |
| cel-miR-5546-3p | down | R03H4.6.1 | up | bus-1 |
| cel-miR-5546-3p | down | F14B8.5b.1 | up | F14B8.5 |
| cel-miR-5546-3p | down | F14B8.5a.1 | up | F14B8.5 |
| cel-miR-5546-3p | down | Y47H9C.10.1 | up | fbxa-216 |
| cel-miR-5546-3p | down | Y43C5A.3.1 | up | Y43C5A.3 |
| cel-miR-5546-3p | down | C05D12.7.1 | up | C05D12.7 |
| cel-miR-5546-3p | down | W02C12.3b.1 | up | hlh-30 |
| cel-miR-5546-3p | down | F22E10.3.1 | up | pgp-14 |
| cel-miR-5546-3p | down | W02F12.2.1 | up | W02F12.2 |
| cel-miR-5546-3p | down | ZK742.1a.3 | up | xpo-1 |
| cel-miR-5546-3p | down | Y22D7AL.8d.1 | up | sms-3 |
| cel-miR-5546-3p | down | W05H7.4b.1 | up | zfp-3 |
| cel-miR-5546-3p | down | Y54G11A.4.1 | up | Y54G11A.4 |
| cel-miR-5546-3p | down | F37C4.5a.2 | up | F37C4.5 |
| cel-miR-5546-3p | down | F37C4.5a.1 | up | F37C4.5 |
| cel-miR-5546-3p | down | F37C4.5a.3 | up | F37C4.5 |
| cel-miR-5546-3p | down | C06E1.8.1 | up | ztf-30 |
| cel-miR-5546-3p | down | F49E11.1c.1 | up | mbk-2 |
| cel-miR-5546-3p | down | H19J13.1.1 | up | otpl-8 |
| cel-miR-5546-3p | down | K10G6.3k.1 | up | sea-2 |
| cel-miR-5546-3p | down | C14B9.4b.1 | up | plk-1 |
| cel-miR-5546-3p | down | B0228.4c.1 | up | cpna-2 |
| cel-miR-5546-3p | down | E02H9.3a.1 | up | E02H9.3 |
| cel-miR-5546-3p | down | C40H1.8.1 | up | C40H1.8 |
| cel-miR-5546-3p | down | Y47H9C.4d.1 | up | ced-1 |
| cel-miR-5546-3p | down | Y80D3A.10.1 | up | nlp-42 |
| cel-miR-5546-3p | down | F53A2.9.1 | up | F53A2.9 |
| cel-miR-5546-3p | down | H06H21.10a.1 | up | tat-2 |
| cel-miR-5546-3p | down | H06H21.10b.1 | up | tat-2 |
| cel-miR-5546-3p | down | F53F4.13.1 | up | F53F4.13 |
| cel-miR-5546-3p | down | C38H2.2.2 | up | C38H2.2 |
| cel-miR-5546-3p | down | C25A11.4c.1 | up | ajm-1 |
| cel-miR-5546-3p | down | ZK524.2d.1 | up | unc-13 |
| cel-miR-5546-3p | down | C54D2.5a.1 | up | cca-1 |
| cel-miR-5546-3p | down | C16D9.4.1 | up | C16D9.4 |
| cel-miR-5546-3p | down | F01G12.5b.1 | up | let-2 |
| cel-miR-5546-3p | down | F52E10.4c.1 | up | F52E10.4 |
| cel-miR-5546-3p | down | F54A5.1.1 | up | hmbx-1 |
| cel-miR-5546-3p | down | Y47D7A.5.1 | up | grl-5 |
| cel-miR-5546-3p | down | F58F9.1a.1 | up | F58F9.1 |
| cel-miR-5546-3p | down | ZK795.1.1 | up | ipmk-1 |
| cel-miR-5546-3p | down | E04A4.7.1 | up | cyc-2.1 |
| cel-miR-5546-3p | down | F15G9.4b.1 | up | him-4 |
| cel-miR-5546-3p | down | ZK783.1k.1 | up | fbn-1 |
| cel-miR-5546-3p | down | ZK783.1b.1 | up | fbn-1 |
| cel-miR-5546-3p | down | ZK783.1f.1 | up | fbn-1 |
| cel-miR-5546-3p | down | C07G2.2c.2 | up | atf-7 |
| cel-miR-5546-3p | down | F13H8.5b.1 | up | F13H8.5 |
| cel-miR-5546-3p | down | C07G2.2d.1 | up | atf-7 |
| cel-miR-5546-3p | down | F56D6.2.2 | up | clec-67 |
| cel-miR-5546-3p | down | ZK836.3.1 | up | ZK836.3 |
| cel-miR-58b-5p | down | F58D5.5.1 | up | F58D5.5 |
| cel-miR-58b-5p | down | F25E5.8b.1 | up | F25E5.8 |
| cel-miR-58b-5p | down | ZK795.1.1 | up | ipmk-1 |
| cel-miR-58b-5p | down | C14C10.3c.1 | up | ril-2 |
| cel-miR-58b-5p | down | F37B12.1.1 | up | F37B12.1 |
| cel-miR-58b-5p | down | Y15E3A.5.1 | up | Y15E3A.5 |
| cel-miR-58b-5p | down | W06F12.1c.1 | up | lit-1 |
| cel-miR-58b-5p | down | Y47G6A.30.1 | up | Y47G6A.30 |
| cel-miR-77-3p | down | C14C10.3c.1 | up | ril-2 |
| cel-miR-77-3p | down | C32F10.8a.1 | up | C32F10.8 |
| cel-miR-77-3p | down | ZK1307.1a.1 | up | ZK1307.1 |
| cel-miR-77-3p | down | F11A10.6a.2 | up | F11A10.6 |
| cel-miR-77-3p | down | C44C10.11.1 | up | C44C10.11 |
| cel-miR-77-3p | down | C44C10.11.2 | up | C44C10.11 |
| cel-miR-77-3p | down | C08F1.4a.3 | up | math-3 |
| cel-miR-77-3p | down | F36H5.3a.1 | up | math-28 |
| cel-miR-77-3p | down | F53F10.2c.4 | up | F53F10.2 |
| cel-miR-77-3p | down | F31F6.4b.1 | up | flp-8 |
| cel-miR-77-3p | down | F57C12.4.1 | up | mrp-2 |
| cel-miR-77-3p | down | M03A1.7.1 | up | dao-2 |
| cel-miR-77-3p | down | M03A1.7.2 | up | dao-2 |
| cel-miR-77-3p | down | C05D12.3c.1 | up | C05D12.3 |
| cel-miR-77-3p | down | C05D12.3c.2 | up | C05D12.3 |
| cel-miR-77-3p | down | F36H5.14.1 | up | F36H5.14 |
| cel-miR-77-3p | down | ZC168.4.3 | up | cyb-1 |
| cel-miR-77-3p | down | C42D8.5a.1 | up | acn-1 |
| cel-miR-77-3p | down | H23N18.3.1 | up | ugt-8 |
| cel-miR-77-3p | down | C47D2.1.1 | up | C47D2.1 |
| cel-miR-77-3p | down | R08F11.4.1 | up | R08F11.4 |
| cel-miR-77-5p | down | C13B9.4d.1 | up | pdfr-1 |
| cel-miR-77-5p | down | F49E2.5h.2 | up | F49E2.5 |
| cel-miR-77-5p | down | C46C2.1n.2 | up | wnk-1 |
| cel-miR-77-5p | down | C46C2.1d.1 | up | wnk-1 |
| cel-miR-77-5p | down | K02E10.2b.1 | up | hid-1 |
| cel-miR-77-5p | down | B0222.5.1 | up | B0222.5 |
| cel-miR-77-5p | down | R13F6.3.1 | up | srv-1 |
| cel-miR-77-5p | down | C34G6.6a.1 | up | noah-1 |
| cel-miR-77-5p | down | C34G6.6c.1 | up | noah-1 |
| cel-miR-77-5p | down | B0284.1.1 | up | pals-26 |
| cel-miR-77-5p | down | M110.5b.2 | up | dab-1 |
| cel-miR-77-5p | down | M110.5c.1 | up | dab-1 |
| cel-miR-77-5p | down | M163.1.1 | up | M163.1 |
| cel-miR-77-5p | down | F49E11.1c.1 | up | mbk-2 |
| cel-miR-77-5p | down | F09A5.4e.2 | up | mob-2 |
| cel-miR-77-5p | down | F59B1.2.1 | up | F59B1.2 |
| cel-miR-77-5p | down | C44C8.6.2 | up | mak-2 |
| cel-miR-77-5p | down | Y75B8A.11a.1 | up | lury-1 |
| cel-miR-77-5p | down | Y71F9B.5b.1 | up | lin-17 |
| cel-miR-77-5p | down | F08B1.1b.1 | up | vhp-1 |
| cel-miR-77-5p | down | F08B1.1e.1 | up | vhp-1 |
| cel-miR-77-5p | down | F08B1.1a.2 | up | vhp-1 |
| cel-miR-77-5p | down | F08B1.1a.1 | up | vhp-1 |
| cel-miR-77-5p | down | F08B1.1f.1 | up | vhp-1 |
| cel-miR-77-5p | down | C18B2.6.1 | up | del-9 |
| cel-miR-77-5p | down | CD4.2.1 | up | crn-2 |
| cel-miR-77-5p | down | F25E5.8b.1 | up | F25E5.8 |
| cel-miR-77-5p | down | F29G9.4a.1 | up | fos-1 |
| cel-miR-77-5p | down | F29G9.4b.1 | up | fos-1 |
| cel-miR-77-5p | down | C31C9.2.1 | up | C31C9.2 |
| cel-miR-784-5p | down | F58B4.1b.1 | up | nas-31 |
| cel-miR-784-5p | down | C52G5.2.1 | up | C52G5.2 |
| cel-miR-784-5p | down | W06F12.1c.1 | up | lit-1 |
| cel-miR-784-5p | down | C54G10.4b.1 | up | slc-25A29 |
| cel-miR-784-5p | down | F35D11.2a.1 | up | pqn-35 |
| cel-miR-784-5p | down | Y75B8A.22.1 | up | tim-1 |
| cel-miR-784-5p | down | C53B4.4d.1 | up | C53B4.4 |
| cel-miR-784-5p | down | C53B4.4g.1 | up | C53B4.4 |
| cel-miR-784-5p | down | B0303.7a.1 | up | B0303.7 |
| cel-miR-784-5p | down | F14B8.5b.1 | up | F14B8.5 |
| cel-miR-784-5p | down | F14B8.5a.1 | up | F14B8.5 |
| cel-miR-784-5p | down | F56D12.5a.3 | up | vig-1 |
| cel-miR-784-5p | down | F56D12.5b.1 | up | vig-1 |
| cel-miR-784-5p | down | F25D1.1a.1 | up | ppm-1.A |
| cel-miR-784-5p | down | F18A1.2a.1 | up | lin-26 |
| cel-miR-784-5p | down | F38E11.7b.1 | up | wrt-3 |
| cel-miR-784-5p | down | ZK418.9b.2 | up | fubl-2 |
| cel-miR-784-5p | down | F56D2.1a.2 | up | ucr-1 |
| cel-miR-784-5p | down | F56D2.1c.1 | up | ucr-1 |
| cel-miR-784-5p | down | Y73F4A.2.1 | up | Y73F4A.2 |
| cel-miR-78 | down | F57B9.6a.3 | up | inf-1 |
| cel-miR-78 | down | C05D12.7.1 | up | C05D12.7 |
| cel-miR-78 | down | K09E2.4b.3 | up | rig-1 |
| cel-miR-78 | down | Y43F8B.1d.1 | up | Y43F8B.1 |
| cel-miR-78 | down | Y43F8B.1a.2 | up | Y43F8B.1 |
| cel-miR-78 | down | B0412.2.1 | up | daf-7 |
| cel-miR-78 | down | Y66H1B.2c.3 | up | fln-1 |
| cel-miR-8200-3p | down | D2023.1f.1 | up | D2023.1 |
| cel-miR-8200-3p | down | C33D9.3b.1 | up | C33D9.3 |
| cel-miR-8200-3p | down | C16A11.5c.1 | up | C16A11.5 |
| cel-miR-8200-3p | down | F22A3.1b.1 | up | ets-4 |
| cel-miR-8200-3p | down | F40F9.2.2 | up | tag-120 |
| cel-miR-8200-3p | down | C14C10.3c.1 | up | ril-2 |
| cel-miR-8200-3p | down | Y48G8AL.13a.2 | up | Y48G8AL.13 |
| cel-miR-8200-3p | down | Y48G8AL.13a.3 | up | Y48G8AL.13 |
| cel-miR-8200-3p | down | Y48G8AL.13b.1 | up | Y48G8AL.13 |
| cel-miR-8200-3p | down | F56D2.1a.2 | up | ucr-1 |
| cel-miR-8200-3p | down | F56D2.1c.1 | up | ucr-1 |
| cel-miR-8200-3p | down | H10E21.5.1 | up | H10E21.5 |
| cel-miR-8200-3p | down | F15A8.6a.1 | up | F15A8.6 |
| cel-miR-8200-3p | down | ZC262.2b.1 | up | ZC262.2 |
| cel-miR-8200-3p | down | F42G2.4.1 | up | fbxa-182 |
| cel-miR-8200-3p | down | F01G10.9.1 | up | F01G10.9 |
| cel-miR-8200-3p | down | K02E10.2b.1 | up | hid-1 |
| cel-miR-8200-3p | down | W09C2.3b.1 | up | mca-1 |
| cel-miR-8200-3p | down | W09C2.3c.1 | up | mca-1 |
| cel-miR-8200-3p | down | K09A9.1.1 | up | nipi-3 |
| cel-miR-8200-3p | down | F18H3.4.1 | up | F18H3.4 |
| cel-miR-8200-3p | down | Y95B8A.1.1 | up | nas-30 |
| cel-miR-8200-3p | down | F55F8.1.1 | up | ptr-10 |
| cel-miR-8200-3p | down | Y9C2UA.1c.1 | up | Y9C2UA.1 |
| cel-miR-8200-3p | down | C56G2.1b.1 | up | akap-1 |

Supplementary Table 4 The prediction of miRNA-mRNA pairs of lncRNA downstream.

| **microRNA** | **direction** | **mRNA** | **direction** | **parent gene** |
| --- | --- | --- | --- | --- |
| cel-miR-4807 | down | R03G5.1c.1 | up | eef-1A.2 |
| cel-miR-4807 | down | F58D5.5.1 | up | F58D5.5 |
| cel-miR-4807 | down | F21A10.2d.1 | up | myrf-2 |
| cel-miR-4807 | down | K11E8.1d.1 | up | unc-43 |
| cel-miR-4807 | down | K11E8.1e.1 | up | unc-43 |
| cel-miR-4807 | down | K11E8.1o.1 | up | unc-43 |
| cel-miR-4807 | down | K11E8.1r.1 | up | unc-43 |
| cel-miR-4807 | down | C25E10.5.1 | up | C25E10.5 |
| cel-miR-4807 | down | Y37A1B.2h.3 | up | lst-4 |
| cel-miR-4807 | down | Y37E3.16.2 | up | fpn-1.1 |
| cel-miR-4807 | down | Y37E3.16.1 | up | fpn-1.1 |
| cel-miR-4807 | down | C44C10.11.1 | up | C44C10.11 |
| cel-miR-4807 | down | F49E11.1c.1 | up | mbk-2 |
| cel-miR-4807 | down | ZK177.8b.2 | up | ZK177.8 |
| cel-miR-4807 | down | C46C2.1n.2 | up | wnk-1 |
| cel-miR-4807 | down | C46C2.1d.1 | up | wnk-1 |
| cel-miR-4807 | down | F30A10.2.1 | up | F30A10.2 |
| cel-miR-4807 | down | F55H12.4.1 | up | F55H12.4 |
| cel-miR-4807 | down | R03G5.1a.3 | up | eef-1A.2 |
| cel-miR-4807 | down | R03G5.1a.2 | up | eef-1A.2 |
| cel-miR-4807 | down | R03G5.1d.2 | up | eef-1A.2 |
| cel-miR-4807 | down | R03G5.1d.1 | up | eef-1A.2 |
| cel-miR-4807 | down | R03G5.1b.1 | up | eef-1A.2 |
| cel-miR-4807 | down | F55A11.4a.2 | up | F55A11.4 |
| cel-miR-5546-3p | down | R166.1a.1 | up | mab-10 |
| cel-miR-5546-3p | down | Y55B1BL.1a.1 | up | Y55B1BL.1 |
| cel-miR-5546-3p | down | K04G11.5c.1 | up | irk-3 |
| cel-miR-5546-3p | down | K04G11.5e.1 | up | irk-3 |
| cel-miR-5546-3p | down | R03H4.6.1 | up | bus-1 |
| cel-miR-5546-3p | down | F14B8.5b.1 | up | F14B8.5 |
| cel-miR-5546-3p | down | F14B8.5a.1 | up | F14B8.5 |
| cel-miR-5546-3p | down | Y47H9C.10.1 | up | fbxa-216 |
| cel-miR-5546-3p | down | Y43C5A.3.1 | up | Y43C5A.3 |
| cel-miR-5546-3p | down | C05D12.7.1 | up | C05D12.7 |
| cel-miR-5546-3p | down | W02C12.3b.1 | up | hlh-30 |
| cel-miR-5546-3p | down | F22E10.3.1 | up | pgp-14 |
| cel-miR-5546-3p | down | W02F12.2.1 | up | W02F12.2 |
| cel-miR-5546-3p | down | ZK742.1a.3 | up | xpo-1 |
| cel-miR-5546-3p | down | Y22D7AL.8d.1 | up | sms-3 |
| cel-miR-5546-3p | down | W05H7.4b.1 | up | zfp-3 |
| cel-miR-5546-3p | down | Y54G11A.4.1 | up | Y54G11A.4 |
| cel-miR-5546-3p | down | F37C4.5a.2 | up | F37C4.5 |
| cel-miR-5546-3p | down | F37C4.5a.1 | up | F37C4.5 |
| cel-miR-5546-3p | down | F37C4.5a.3 | up | F37C4.5 |
| cel-miR-5546-3p | down | C06E1.8.1 | up | ztf-30 |
| cel-miR-5546-3p | down | F49E11.1c.1 | up | mbk-2 |
| cel-miR-5546-3p | down | H19J13.1.1 | up | otpl-8 |
| cel-miR-5546-3p | down | K10G6.3k.1 | up | sea-2 |
| cel-miR-5546-3p | down | C14B9.4b.1 | up | plk-1 |
| cel-miR-5546-3p | down | B0228.4c.1 | up | cpna-2 |
| cel-miR-5546-3p | down | E02H9.3a.1 | up | E02H9.3 |
| cel-miR-5546-3p | down | C40H1.8.1 | up | C40H1.8 |
| cel-miR-5546-3p | down | Y47H9C.4d.1 | up | ced-1 |
| cel-miR-5546-3p | down | Y80D3A.10.1 | up | nlp-42 |
| cel-miR-5546-3p | down | F53A2.9.1 | up | F53A2.9 |
| cel-miR-5546-3p | down | H06H21.10a.1 | up | tat-2 |
| cel-miR-5546-3p | down | H06H21.10b.1 | up | tat-2 |
| cel-miR-5546-3p | down | F53F4.13.1 | up | F53F4.13 |
| cel-miR-5546-3p | down | C38H2.2.2 | up | C38H2.2 |
| cel-miR-5546-3p | down | C25A11.4c.1 | up | ajm-1 |
| cel-miR-5546-3p | down | ZK524.2d.1 | up | unc-13 |
| cel-miR-5546-3p | down | C54D2.5a.1 | up | cca-1 |
| cel-miR-5546-3p | down | C16D9.4.1 | up | C16D9.4 |
| cel-miR-5546-3p | down | F01G12.5b.1 | up | let-2 |
| cel-miR-5546-3p | down | F52E10.4c.1 | up | F52E10.4 |
| cel-miR-5546-3p | down | F54A5.1.1 | up | hmbx-1 |
| cel-miR-5546-3p | down | Y47D7A.5.1 | up | grl-5 |
| cel-miR-5546-3p | down | F58F9.1a.1 | up | F58F9.1 |
| cel-miR-5546-3p | down | ZK795.1.1 | up | ipmk-1 |
| cel-miR-5546-3p | down | E04A4.7.1 | up | cyc-2.1 |
| cel-miR-5546-3p | down | F15G9.4b.1 | up | him-4 |
| cel-miR-5546-3p | down | ZK783.1k.1 | up | fbn-1 |
| cel-miR-5546-3p | down | ZK783.1b.1 | up | fbn-1 |
| cel-miR-5546-3p | down | ZK783.1f.1 | up | fbn-1 |
| cel-miR-5546-3p | down | C07G2.2c.2 | up | atf-7 |
| cel-miR-5546-3p | down | F13H8.5b.1 | up | F13H8.5 |
| cel-miR-5546-3p | down | C07G2.2d.1 | up | atf-7 |
| cel-miR-5546-3p | down | F56D6.2.2 | up | clec-67 |
| cel-miR-5546-3p | down | ZK836.3.1 | up | ZK836.3 |
| cel-miR-58b-5p | down | F58D5.5.1 | up | F58D5.5 |
| cel-miR-58b-5p | down | F25E5.8b.1 | up | F25E5.8 |
| cel-miR-58b-5p | down | ZK795.1.1 | up | ipmk-1 |
| cel-miR-58b-5p | down | C14C10.3c.1 | up | ril-2 |
| cel-miR-58b-5p | down | F37B12.1.1 | up | F37B12.1 |
| cel-miR-58b-5p | down | Y15E3A.5.1 | up | Y15E3A.5 |
| cel-miR-58b-5p | down | W06F12.1c.1 | up | lit-1 |
| cel-miR-58b-5p | down | Y47G6A.30.1 | up | Y47G6A.30 |
| cel-miR-784-5p | down | F58B4.1b.1 | up | nas-31 |
| cel-miR-784-5p | down | C52G5.2.1 | up | C52G5.2 |
| cel-miR-784-5p | down | W06F12.1c.1 | up | lit-1 |
| cel-miR-784-5p | down | C54G10.4b.1 | up | slc-25A29 |
| cel-miR-784-5p | down | F35D11.2a.1 | up | pqn-35 |
| cel-miR-784-5p | down | Y75B8A.22.1 | up | tim-1 |
| cel-miR-784-5p | down | C53B4.4d.1 | up | C53B4.4 |
| cel-miR-784-5p | down | C53B4.4g.1 | up | C53B4.4 |
| cel-miR-784-5p | down | B0303.7a.1 | up | B0303.7 |
| cel-miR-784-5p | down | F14B8.5b.1 | up | F14B8.5 |
| cel-miR-784-5p | down | F14B8.5a.1 | up | F14B8.5 |
| cel-miR-784-5p | down | F56D12.5a.3 | up | vig-1 |
| cel-miR-784-5p | down | F56D12.5b.1 | up | vig-1 |
| cel-miR-784-5p | down | F25D1.1a.1 | up | ppm-1.A |
| cel-miR-784-5p | down | F18A1.2a.1 | up | lin-26 |
| cel-miR-784-5p | down | F38E11.7b.1 | up | wrt-3 |
| cel-miR-784-5p | down | ZK418.9b.2 | up | fubl-2 |
| cel-miR-784-5p | down | F56D2.1a.2 | up | ucr-1 |
| cel-miR-784-5p | down | F56D2.1c.1 | up | ucr-1 |
| cel-miR-784-5p | down | Y73F4A.2.1 | up | Y73F4A.2 |
| cel-miR-789-2-5p | down | Y54G11A.4.1 | up | Y54G11A.4 |
| cel-miR-789-2-5p | down | B0403.3.1 | up | B0403.3 |
| cel-miR-789-2-5p | down | F13B12.5.1 | up | ins-1 |
| cel-miR-789-2-5p | down | F35B12.6.1 | up | tag-290 |
| cel-miR-789-2-5p | down | F55A11.4a.2 | up | F55A11.4 |
| cel-miR-789-2-5p | down | R13H8.1e.1 | up | daf-16 |
| cel-miR-789-2-5p | down | Y39A3CL.5a.1 | up | clp-4 |
| cel-miR-789-2-5p | down | Y22D7AL.8d.1 | up | sms-3 |
| cel-miR-789-2-5p | down | F09C3.2.1 | up | F09C3.2 |
| cel-miR-789-2-5p | down | Y48G1BR.1.1 | up | Y48G1BR.1 |
| cel-miR-789-2-5p | down | B0464.7.1 | up | baf-1 |
| cel-miR-789-2-5p | down | R13H8.1b.2 | up | daf-16 |
| cel-miR-789-2-5p | down | C02F12.1b.1 | up | tsp-17 |
| cel-miR-789-2-5p | down | W04G3.3.1 | up | lpr-4 |
| cel-miR-789-2-5p | down | K07C11.7b.1 | up | K07C11.7 |
| cel-miR-789-2-5p | down | ZK270.2b.1 | up | frm-1 |
| cel-miR-789-2-5p | down | R06C7.5b.1 | up | adsl-1 |
| cel-miR-789-2-5p | down | C41D11.1b.1 | up | eri-6 |
| cel-miR-789-2-5p | down | C01F6.6i.1 | up | nrfl-1 |
| cel-miR-789-2-5p | down | C01F6.6b.1 | up | nrfl-1 |
| cel-miR-789-2-5p | down | C01F6.6e.4 | up | nrfl-1 |
| cel-miR-789-2-5p | down | C37A2.5b.2 | up | pqn-21 |
| cel-miR-789-2-5p | down | K09A9.1.1 | up | nipi-3 |
| cel-miR-789-2-5p | down | Y15E3A.1a.3 | up | nhr-91 |
| cel-miR-789-2-5p | down | R02F2.1a.3 | up | R02F2.1 |
| cel-miR-789-2-5p | down | R02F2.1b.3 | up | R02F2.1 |
| cel-miR-789-2-5p | down | E03G2.4.1 | up | col-186 |
| cel-miR-789-2-5p | down | C40C9.5b.1 | up | nlg-1 |
| cel-miR-789-2-5p | down | D1005.3.2 | up | cebp-1 |
| cel-miR-789-2-5p | down | D1005.3.1 | up | cebp-1 |
| cel-miR-789-2-5p | down | F14D12.2.1 | up | unc-97 |
| cel-miR-789-2-5p | down | F58B3.5c.1 | up | mars-1 |
| cel-miR-789-2-5p | down | Y16E11A.1.1 | up | math-44 |
| cel-miR-789-2-5p | down | F19B10.1.2 | up | F19B10.1 |
| cel-miR-789-2-5p | down | F31E3.2b.1 | up | F31E3.2 |
| cel-miR-789-2-5p | down | ZK520.2.1 | up | sid-2 |
| cel-miR-789-2-5p | down | F31F6.6.1 | up | nac-1 |
| cel-miR-789-2-5p | down | K09E2.4b.3 | up | rig-1 |
| cel-miR-8196b-3p | down | H19N07.2a.1 | up | math-33 |
| cel-miR-8196b-3p | down | H19N07.2b.1 | up | math-33 |
| cel-miR-8196b-3p | down | F46G10.4.1 | up | F46G10.4 |
| cel-miR-8196b-3p | down | B0507.10.1 | up | ddn-1 |
| cel-miR-8196b-3p | down | Y17G7B.20e.1 | up | Y17G7B.20 |
| cel-miR-8196b-3p | down | Y17G7B.20c.2 | up | Y17G7B.20 |
| cel-miR-8196b-3p | down | Y17G7B.20b.2 | up | Y17G7B.20 |
| cel-miR-8196b-3p | down | Y17G7B.20a.1 | up | Y17G7B.20 |

Supplementary Table 5 The terms associated with production, aging process and nervous system in the results of Go analysis on lncRNA.

| ID | Description | GeneRatio | BgRatio | pvalue | p.adjust | qvalue | geneID | Count |
| --- | --- | --- | --- | --- | --- | --- | --- | --- |
| GO:0009792 | embryo development ending in birth or egg hatching | 6/71 | 305/9624 | 0.024664 | 0.19384 | 0.166641 | mbk-2/ajm-1/let-2/lit-1/tim-1/baf-1 | 6 |
| GO:0009798 | axis specification | 4/71 | 72/9624 | 0.001918 | 0.101961 | 0.087654 | mbk-2/ced-1/lit-1/math-33 | 4 |
| GO:0001505 | regulation of neurotransmitter levels | 3/71 | 73/9624 | 0.01656 | 0.174476 | 0.149995 | unc-13/tsp-17/nlg-1 | 3 |
| GO:0006836 | neurotransmitter transport | 3/71 | 80/9624 | 0.021093 | 0.177036 | 0.152195 | unc-13/tsp-17/nlg-1 | 3 |
| GO:0008286 | insulin receptor signaling pathway | 2/71 | 20/9624 | 0.009357 | 0.140808 | 0.121051 | ins-1/daf-16 | 2 |
| GO:0034599 | cellular response to oxidative stress | 2/71 | 23/9624 | 0.012284 | 0.15666 | 0.134678 | daf-16/tsp-17 | 2 |
| GO:0032868 | response to insulin | 2/71 | 24/9624 | 0.013337 | 0.159722 | 0.137311 | ins-1/daf-16 | 2 |
| GO:0032869 | cellular response to insulin stimulus | 2/71 | 24/9624 | 0.013337 | 0.159722 | 0.137311 | ins-1/daf-16 | 2 |
| GO:0008595 | anterior/posterior axis specification, embryo | 2/71 | 30/9624 | 0.020433 | 0.174476 | 0.149995 | mbk-2/math-33 | 2 |
| GO:0051588 | regulation of neurotransmitter transport | 2/71 | 30/9624 | 0.020433 | 0.174476 | 0.149995 | tsp-17/nlg-1 | 2 |
| GO:0000578 | embryonic axis specification | 2/71 | 32/9624 | 0.023079 | 0.187302 | 0.161021 | mbk-2/math-33 | 2 |
| GO:0110037 | regulation of nematode male tail tip morphogenesis | 2/71 | 39/9624 | 0.033361 | 0.222862 | 0.191591 | plk-1/lit-1 | 2 |

Supplementary Table 6 The predictive result of the kinase-substrate interaction using iGPS.

| Phosphoprotein | Gene name | Kinase ID | Kinase name | Peptide | Position |
| --- | --- | --- | --- | --- | --- |
| H2KYC5 | ajm-1 | Q19266 | pkc-3 | DRKAQELpSEREMREK | 670 |
| P91198 | ppfr-2 | P34556 | cdk-1 | ASSPKAApSPVSSPKS | 320 |
|  |  | O61847 | cdk-2 |  |  |
| Q95Q95 | let-363 | Q17941 | akt-1 | PPVTRATpSPPPPAQK | 1969 |
|  |  | Q9XTG7 | akt-2 |  |  |
|  |  | Q9TW45 | par-1 |  |  |
|  |  | Q9TXI1 | F23C8.8 |  |  |

Supplementary Table 7 The specific binding site between TFs and kinases in prediction of Jaspar.

| Name | Direction | Score | Relative score | Sequence ID | Start | End | Strand | Predicted sequence |
| --- | --- | --- | --- | --- | --- | --- | --- | --- |
| vab-7 | downregulation | 9.25982 | 0.9118316 | >akt-1==V+ | 165 | 172 | + | ATAATGAA |
| ceh-48 | none | 6.18056 | 0.8579261 | >akt-1==V+ | 40 | 48 | + | TGTCAATTT |
| lim-4 | none | 6.57519 | 0.8533672 | >akt-1==V+ | 164 | 172 | + | CATAATGAA |
| vab-7 | downregulation | 6.48518 | 0.8508565 | >akt-1==V+ | 129 | 136 | + | CGAATTAA |
| zfh-2 | none | 11.1226 | 0.913692 | >cdk-2==I+ | 185 | 195 | + | TCTTAATTATC |
| efl-1 | none | 12.8896 | 0.9127394 | >cdk-2==I+ | 112 | 126 | + | GTAGCGCGGGAAAAC |
| lim-4 | none | 8.89205 | 0.909502 | >cdk-2==I+ | 186 | 194 | + | CTTAATTAT |
| vab-7 | downregulation | 7.01337 | 0.862464 | >cdk-2==I+ | 187 | 194 | + | TTAATTAT |
| elt-3 | upregulation | 13.3768 | 0.9746855 | >cdk-1==III+ | 20 | 27 | + | TTTTATCA |
| unc-86 | downregulation | 11.0943 | 0.9396151 | >cdk-1==III+ | 27 | 34 | + | ATATGAAT |
| pal-1 | downregulation | 7.57865 | 0.8806272 | >cdk-1==III+ | 79 | 86 | + | GGAATAAA |
| cebp-1 | upregulation | 11.379 | 0.8612013 | >cdk-1==III+ | 74 | 85 | + | TTTTCGGAATAA |
| fos-1 | upregulation | 7.26009 | 0.8571429 | >cdk-1==III+ | 165 | 171 | + | TGATTCA |
| nhr-6 | none | 7.55321 | 0.8542776 | >cdk-1==III+ | 85 | 92 | + | AATTGTCA |
